# PCNA has specific functions in regulation of metabolism in haematological cells

**DOI:** 10.1101/2020.04.29.067512

**Authors:** Lisa M. Røst, Camilla Olaisen, Animesh Sharma, Aina Nedal, Voin Petrovic, Hans Fredrik N. Kvitvang, Per Bruheim, Marit Otterlei

## Abstract

PCNA’s essential roles in DNA replication and repair are well established, while its recently discovered cytosolic roles are less explored. Here we show that impairing PCNA’s cytosolic scaffold functions led to massive changes in cellular signaling, and most strikingly, a strong reduction in glycolytic metabolite and nucleoside phosphate pools in haematological cancer cells. This was not detected in cells from solid tissues nor in primary monocytes from healthy donors. However, lipopolysaccharide stimulated monocytes responded to targeting PCNA’s scaffold function similarly to haematological cancer cells, suggesting that cellular stress is an important factor for the observed response. Integrated transcriptome, proteome and metabolome analysis revealed that pathways involved in protein stability/folding and immune responses were affected in haematopoietic cancer cells. Altogether, our data suggests that PCNA has an important role in regulation of cytoplasmic stress via regulation of central carbon metabolism in haematological cells, which potentially can be targeted in cancer treatment.

**Highlight:** A strong reduction in glycolytic metabolites and the nucleoside phosphate pools was detected in haematological cancer cells upon inhibiting PCNA scaffold functions. These metabolic changes were not found in cancer cells from other tissues, indicating a special role of PCNA in regulation of metabolism in cells of haematological origin.

## Introduction

Proliferating cell nuclear antigen (PCNA), a member of the essential and highly conserved DNA sliding clamp family, interacts with proteins carrying either of the conserved motifs PIP-box (PCNA interacting peptide – box) (Warbrick, 1998) or APIM (AlkB homolog 2 interacting motif) (Gilljam et al., 2009). These motifs have an overlapping binding site on PCNA (Müller et al., 2013; Sebesta, Cooper, Ariza, Carnie, & Ahel, 2017) but while replication is strictly dependent upon PIP-box - PCNA interactions, data supports an unique role of the APIM - PCNA interaction under cellular stress (Gilljam et al., 2009; Müller et al., 2013; Sogaard, Blindheim, et al., 2018; Sogaard, Moestue, et al., 2018; Sogaard et al., 2019; Warbrick, 2006). Many of the APIM- or PIP-containing proteins are not directly involved in processing of DNA, but have important roles in cellular signaling, e.g. in the PI3K/AKT and MAPK pathways (Olaisen et al., 2018; Olaisen, Müller, Nedal, & Otterlei, 2015). Multi-layered regulatory mechanisms are required to control the repertoire of up to 600 proteins (Olaisen et al., 2018) that may interact with PCNA through these two motifs at any given time. These regulatory mechanisms include affinity driven competition, at least partly mediated via posttranslational modifications (PTMs) of PCNA or PCNA binding proteins, and/or the context of the PCNA complexes (e.g. repair vs replication, nuclear vs cytosolic localization, stages in cell cycle, as well as the presence or absence of DNA) (Mailand, Gibbs-Seymour, & Bekker-Jensen, 2013).

Several lines of evidence support cytosolic or non-canonical roles for PCNA: I) Cytosolic PCNA is reported to regulate apoptosis by binding to procaspases in mature neutrophils (Witko-Sarsat et al., 2010), multiple myeloma (MM) cells (Müller et al., 2013), and neuroblastoma cells (Yin et al., 2015); II) Inhibition of the ability of APIM-containing proteins to interact with PCNA by treating cells with a cell-penetrating peptide containing the APIM sequence (APIM-peptide) strongly reduced the secretion of cytokines from human monocytes stimulated with Toll-like receptor ligands (Olaisen et al., 2015); III) Cytosolic PCNA in cancer cells inhibits activation of natural killer cells, thereby helping cancer cells to evade the immune system (Rosental et al., 2011); IV) Glycolytic enzymes associate with PCNA, and nuclear export of PCNA correlated with increased Warburg-effect in acute myeloid leukaemia (AML) cells (Naryzhny & Lee, 2010; Ohayon et al., 2016); V) Multiple proteins involved in PI3K/AKT and MAPK signaling were found in PCNA pull-downs (Olaisen et al., 2015), and targeting PCNA has been linked to downregulation of EGFR and PI3K/AKT pathways in bladder cancer cells and monocytes (Olaisen et al., 2015; Sogaard, Blindheim, et al., 2018); VI) Cytosolic PCNA participates in reactive oxygen species (ROS) regulation via interaction with a subunit of the NADPH oxidase complex, p47phox (Ohayon et al., 2019); VII) Combination treatments with APIM-peptide and receptor tyrosine kinase inhibitors against EGFR/HER2/VEGFR and cMET showed increased anticancer activity in absence of any DNA-damaging agent (Sogaard et al., 2019); and VIII) A functional conservation of PCNA as a scaffold protein in MAPK signaling has been shown between mammalians and yeast (Olaisen et al., 2018). These cytosolic scaffold functions of PCNA, as well as the verified APIM-PCNA interactions important in DNA damage repair and/or tolerance, are illustrated in Figure 1.

**Figure 1:**
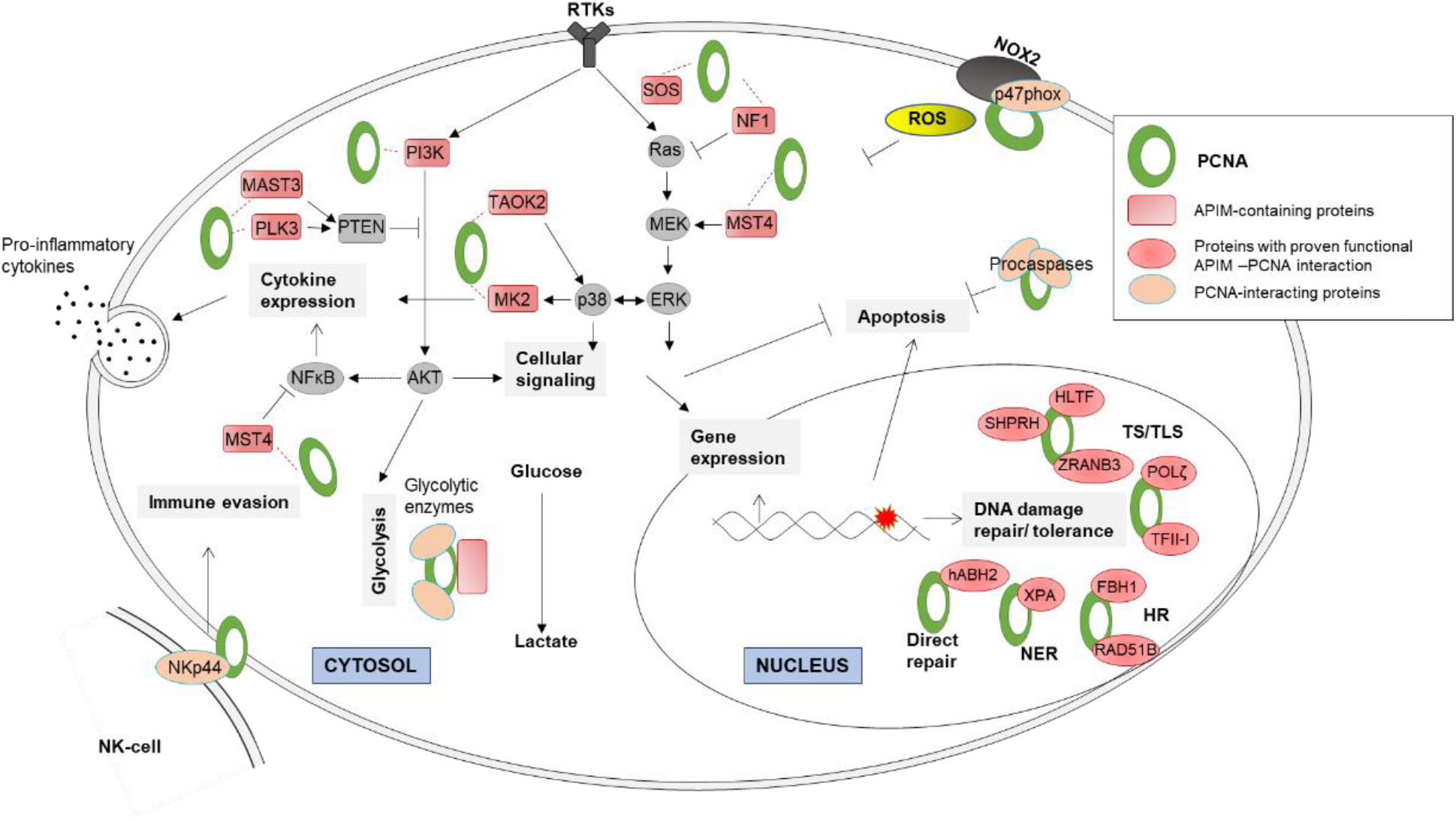
Scaffold functions of PCNA important during cellular stress responses. Nuclear roles: Multiple APIM-containing proteins involved in DNA repair and DNA damage tolerance are shown to be dependent upon their APIM-PCNA interactions for optimal function (Bacquin et al., 2013; Fattah et al., 2014; Gilljam et al., 2009; Gilljam, Müller, Liabakk, & Otterlei, 2012; Raeder et al., 2018; Seelinger & Otterlei, 2020). Cytosolic roles: Multiple cytosolic proteins involved in cellular signaling and metabolism, including glycolytic enzymes, contain APIM or PIP-box motifs (Ohayon et al., 2019; Olaisen et al., 2018; Olaisen et al., 2015), or are found in complex with PCNA (Naryzhny & Lee, 2010; Olaisen et al., 2015; Witko-Sarsat et al., 2010). Studies have shown that impairing PCNA’s scaffold function affects multiple signaling pathways including the PI3K/AKT and MAPK pathways and the cytokine production after toll receptor stimuli (Olaisen et al., 2018; Olaisen et al., 2015; Sogaard et al., 2019), regulation of apoptosis (Müller et al., 2013; Witko-Sarsat et al., 2010; Yin et al., 2015), cytotoxicity of NK cells (Rosental et al., 2011; Shemesh et al., 2018), and cellular defense against ROS (Ohayon et al., 2019).

The affinity of APIM to PCNA is increased during cellular stress (Ciccia et al., 2012; Gilljam et al., 2009) in accordance with the ability of the APIM-peptide to increase the efficacy of multiple anti-cancer drugs, including genotoxic drugs and drugs targeting microtubules or kinases (Gederaas et al., 2014; Müller et al., 2013; Olaisen et al., 2018; Olaisen et al., 2015; Sogaard, Blindheim, et al., 2018; Sogaard, Moestue, et al., 2018; Sogaard et al., 2019). Further, data supports impairment of PCNA’s scaffold roles in DNA repair, DNA translesion synthesis and cellular signaling after APIM-peptide treatment (Gilljam et al., 2012; Olaisen et al., 2015; Raeder et al., 2018). Normal cells have low sensitivity to the APIM-peptide as a single agent, while sensitivity varies between cancer cell lines with haematological cancer cells being most sensitive (Müller et al., 2013). The sensitivity of cancer cells towards the APIM-peptide as a single agent is independent of both their PCNA expression levels and the ratio of PCNA in the cytosol versus the nuclei (Gederaas et al., 2014). Furthermore, apoptosis is induced in all phases of the cell cycle in absence of cell cycle arrest (Müller et al., 2013; Sogaard, Moestue, et al., 2018). This suggests that inhibition of replication is not the primary cause of the anti-cancer effects observed by the APIM-peptide *in vitro* and in several preclinical animal models. This is in accordance with the good tolerability of the APIM-peptide ATX-101 observed in the currently ongoing dose escalating clinical Phase I in cancer patients (all-comers) (ANZCTR:12618001070224), in which prolonged weekly administration in several patients did not reveal any myelosuppressive properties.

Apoptosis is rapidly induced in multiple haematological cancer cell lines, including chronic myeloid leukaemia, T-cell acute lymphoblastic leukaemia, MM, in addition to primary MM cells, after exposure to low doses of APIM-peptide (Müller et al., 2013). The same doses do not induce apoptosis in primary monocytes, but strongly reduce Toll receptor ligand-induced cytokine production (Müller et al., 2013; Olaisen et al., 2015). We have previously also linked PCNA to regulation of PI3K/AKT/mTOR signaling (Olaisen et al., 2018; Olaisen et al., 2015). This pathway is important for regulation of glycolysis, and is often enhanced in cancers of haematological origins such as MM and AML (El Arfani, De Veirman, Maes, De Bruyne, & Menu, 2018; Poulain et al., 2017). This led us to explore whether targeting PCNA with the APIM-peptide affected metabolism, and if there was a difference between the metabolic responses in cells of haematological versus solid tumor origins. Experimental studies of multifunctional proteins and/or proteins with numerous interactions partners, such as PCNA, are challenging. Mutations of the APIM – PCNA interaction site on PCNA is not compatible with life as this is completely overlapping with the PIP-box interactions site, and furthermore, mutations of APIM motifs in ∼100 APIM containing proteins involved in cellular signaling and metabolism is an endless endeavor. Also, robust quantitative methodology at the metabolome and protein systems levels is needed as single metabolite and protein analyses will never be able to reveal the multiple roles of PCNA. Thus, to explore this, we have applied cell viability, metabolome, proteome and transcriptome profiling assays, and ^13^C-label experimentation. Here we show that targeting PCNA with the APIM-peptide reduces metabolite pools of glycolysis, the pentose phosphate pathway (PPP), and nucleoside phosphates in haematological cancer cells and lipopolysaccharide (LPS)-stimulated monocytes, but not in cells from solid tissues. Our findings support a specific and central role of PCNA in regulation of central carbon metabolism in stressed haematological cells, either directly though interaction with glycolytic enzymes or indirectly through interaction with signal network proteins.

## Results

### Targeting PCNA with the APIM-peptide reduced glucose consumption in haematological cancer cells

When examining uptake and excretion fluxes, a reduction in specific glucose consumption and/or lactate excretion rates was detected in four out of five haematological cancer cell lines treated with the APIM-peptide (Figure 2A, E, F, H). Importantly, a mutated APIM-peptide variant (mutAPIM-peptide), with reduced affinity for PCNA (Müller et al., 2013; Olaisen et al., 2015), and a peptide containing only the cell-penetrating part of the APIM-peptide (R11-peptide), resulted in significantly lower and no changes in these rates, respectively, compared to the APIM-peptide treatment (Figure 2A). This shows that the effect of the APIM-peptide is dependent on its affinity to PCNA. Small non-significant reductions were also detected in the embryonic kidney cell line (Hek293, Figure 2C), while increased rates were detected in the prostate cancer cells (DU145, Figure 2D). One acute AML cell line had reduced glucose consumption and lactate production rates after the treatment, but another did not respond at all (NB4 and HL60, Figure 2H and G, respectively).

**Figure 2.**
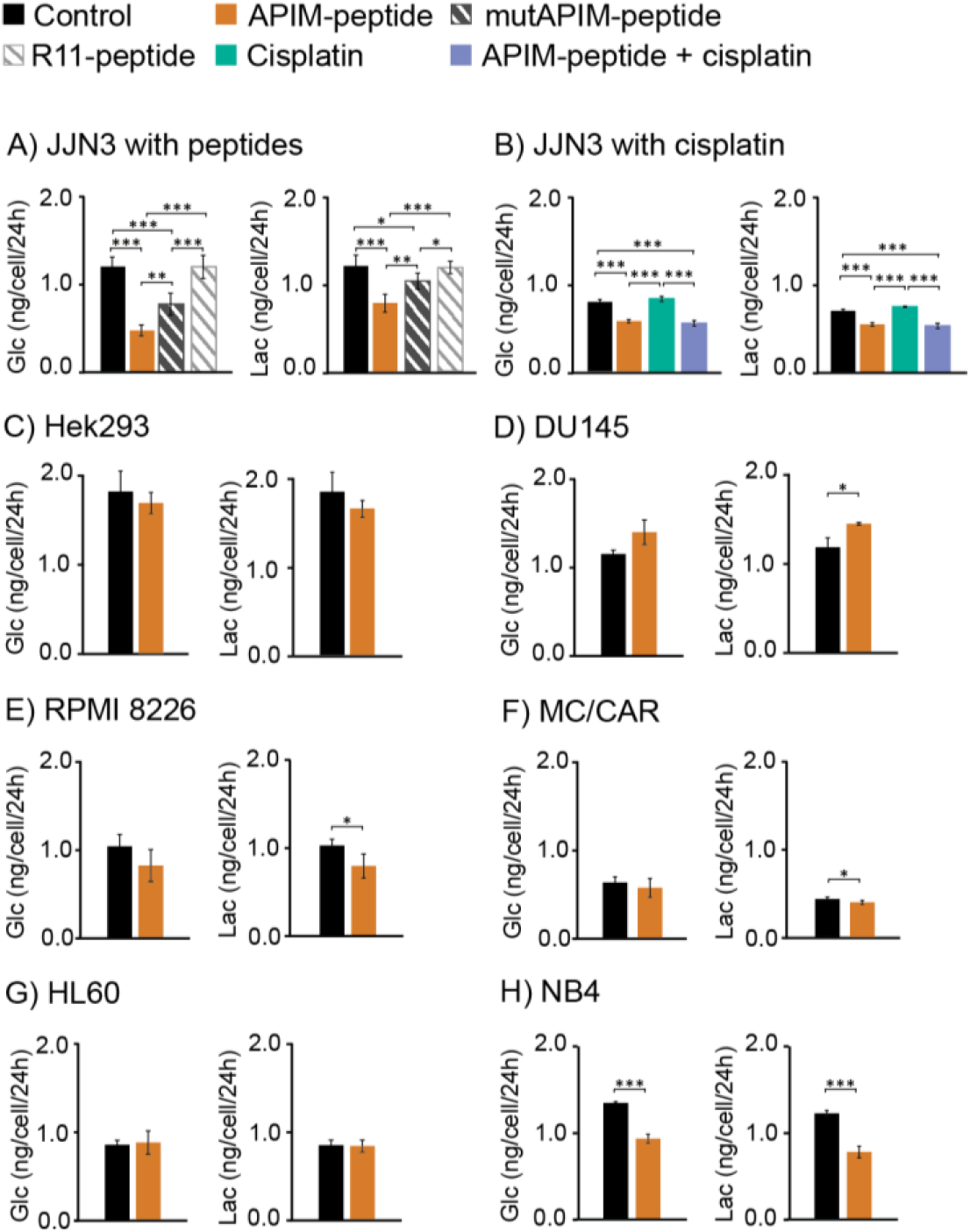
Targeting PCNA with APIM-peptide reduce glucose consumption and lactate excretion in haematological cancer cells. Average glucose (Glc) consumption and lactate (Lac) excretion per cell per 24h ± SD in (**A**) JJN3 cells (MM) treated with APIM-peptide, mutAPIM-peptide, or R11-peptide (8 µM) and (**B**) JJN3 cells treated with APIM-peptide (8 µM), cisplatin (1 µM), or the combination. Significant (*p<0.05, **p<0.01, ***p<0.001) differences (ANOVA, post hoc Tukey’s test) are indicated, n ≥ 4. Average Glc consumption and Lac excretion per cell per 24h ± SD treated with APIM-peptide in (**C**) Hek293 (embryonic kidney, 10 µM), (**D**) DU145 (prostate cancer, 8 µM), (**E**) RPMI 8226 (MM, 8 µM), (**F**) MC/CAR (MM, 8 µM), (**G**) HL60 (AML, 8 µM) and (**H**) NB4 (AML, 8 µM). Significant (*p<0.05, **p<0.01, ***p<0.001) differences (Unpaired two-tailed student T-test) are indicated, n ≥ 3. Details on replicate numbers are listed in Supplementary Table S2.

Including cisplatin as an additional stressor in the treatment regime did not amplify the effects observed in JJN3 (MM), even though the combination of APIM-peptide with cisplatin reduced the growth compared to single treatments after day 2 (Figure 2B)(Supplementary Figure S1). Growth of DU145 cells are also inhibited by the APIM-peptide, still the glucose consumption increased. In Hek293 cells which are not growth-inhibited by the APIM-peptide, the glucose consumption was reduced (Supplementary Figure S1). Thus, changes in glucose uptake rates measured in these experiments are not directly linked to proliferation.

No definite conclusions regarding differences between tumors from solid tissue and haematological cells can be drawn based on uptake and excretion fluxes of this limited panel of cell lines; however, a clear tendency towards a reduction in glucose consumption upon targeting PCNA with the APIM-peptide was detected in the haematological cancer cells.

### Central carbon metabolite pools in haematological cancer cells decrease dramatically when the scaffold function of PCNA is impaired

Further, we extended the panel of cells and compared the effect of targeting PCNA on central metabolism in five haematological cancer cell lines (3x MM, 2x AML), primary monocytes from three healthy donors and three cell lines from solid tissue; kidney, prostate and bladder. Three targeted quantitative liquid chromatography (LC)/capillary ion chromatography (capIC) tandem mass spectrometry (MS/MS) methods, all employing ^13^C-isotope internal standards for absolute quantifications of intracellular concentrations were applied. The data reveals that haematological cancer cell lines responded with a strong reduction in multiple metabolite pools, while smaller changes, or even a modest increase was detected in primary monocytes and cells from solid tissue after treatment with the APIM-peptide (Figure 3A, details in Supplementary Table S1B and in (Sogaard, Blindheim, et al., 2018)).

**Figure 3:**
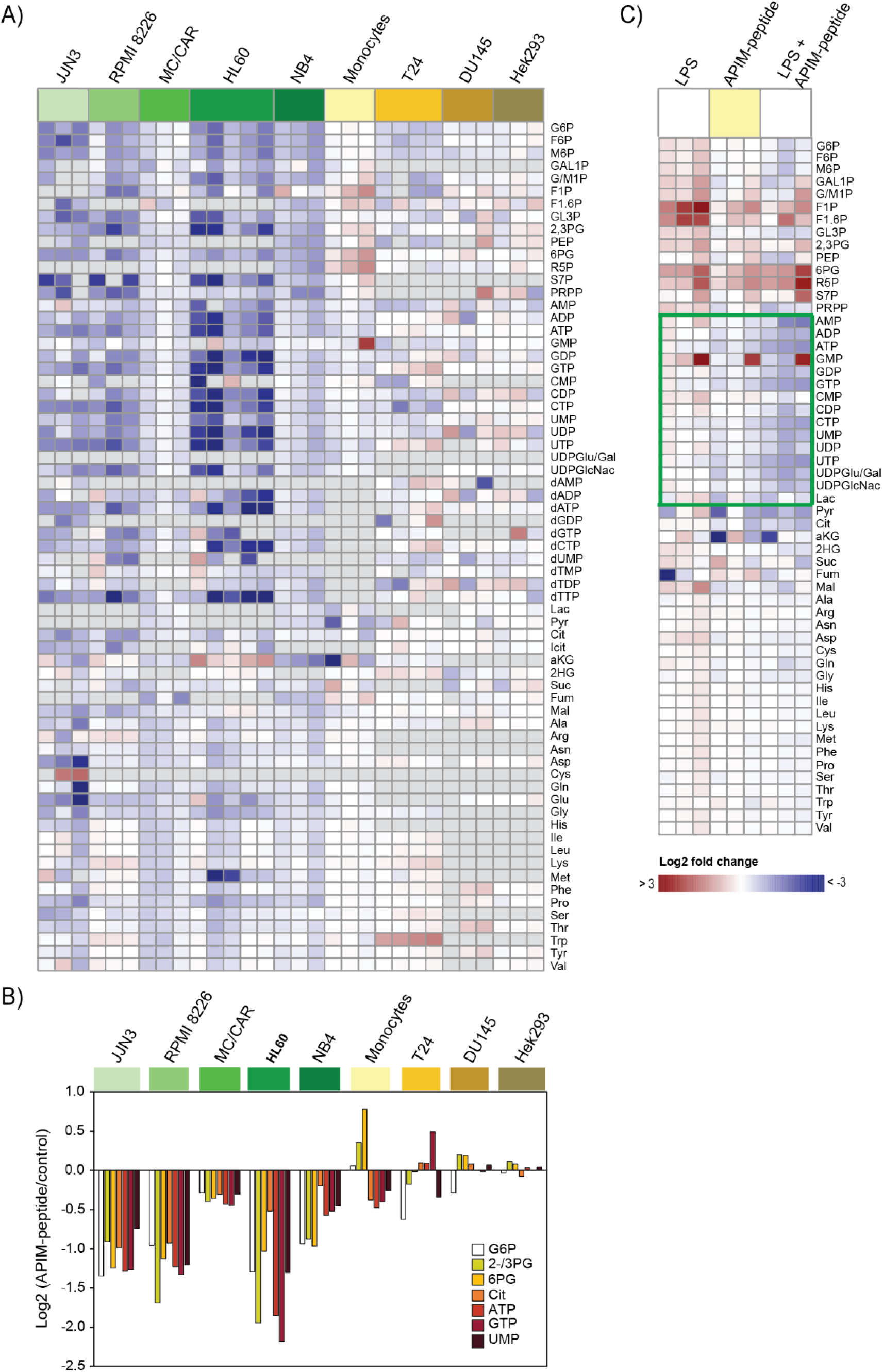
The metabolite profiles of haematological cancer cells is strongly affected by targeting PCNA, whilst the metabolite profiles of primary monocytes and cell lines of solid tissue are not. Log_2_ fold change of all quantified central carbon metabolite pools in APIM-peptide treated (8 µM) cells relative to untreated control after 4h (**A**) heat mapped for JJN3, RPMI 8226, and MC/CAR (MM), HL60 and NB4 (AML), (all variants of green), primary monocytes (yellow), T24 (bladder, 16 µM APIM-peptide, 24h)(light orange), DU145 (prostate 4, 8, 24h) (light brown) and Hek293 (embryonic kidney, 4, 8, 24h) (brown), and (**B**) graphed for metabolites significantly differing (p<0.05, ANOVA, post-hoc Tukey’s test) in ≤ 30% of all possible pairwise comparisons between haematological cancer cells and primary monocytes or cancers cells of solid tissue. (**C**) Heat mapped log_2_ fold change of all measured central carbon metabolite pools in primary monocytes from three donors, listed for LPS (10 ng/ml), APIM-peptide (8µM) (yellow, same data as monocytes in A and B) and the combination relative to untreated control. Grey colour in heat map = NA. Each of n ≥ 3 from one (RPMI 8226, HL60, T24, DU145, and Hek293), or the average of n ≥ 3 from three (JJN3, MC/CAR, NB4, monocytes) repeated experiments is shown. Details on replicate numbers are listed in Supplementary Table S2, and metabolite abbreviations are listed in Supplementary Table S1A.

Out of the metabolites measured in all cell types, the levels of ten metabolites were significantly different (p<0.05) between haematological cancer cell lines and primary monocytes or cell lines from solid tissue in >30% of all possible pairwise comparisons (Supplementary Table S1C). Seven out of these metabolites, all having important roles in central carbon metabolism; G6P, 2-/3PG, 6PG, Cit, ATP, GTP and UMP, are depicted in Figure 3B (for metabolite abbreviations see Supplementary Table S1A). Notably, primary monocytes responded like the haematological cancer cells with respect to reduction in the nucleoside phosphate pools but have a more similar response as the cells from solid tissue for the other central metabolite pools (glycolysis, PPP and TCA) (Figure 3A and B). On the other hand, stimulation of the monocytes with the toll-like receptor 4 ligand LPS, resulted in increased nucleoside phosphate pools and glycolytic intermediates (Figure 3C). However, when monocytes were treated with a combination of LPS and APIM-peptide, most metabolite pools were reduced (Figure 3C). This trend is most prominent for the nucleoside phosphate pools (framed in Figure 3C) where the combination treatment results in downregulation like the other haematological cancer cells treated with APIM-peptide (green panels in Figure 3A). Previous data has shown increased affinity for APIM – PCNA interactions during cellular stress (Ciccia et al., 2012; Gilljam et al., 2009). Therefore, this strong reduction in the nucleoside phosphate pools may suggest that stress induced in the monocytes by LPS increased the affinity of the APIM-peptide to PCNA and thereby the effect of the APIM-peptide treatment. Replicative stress is often elevated in cancer cells; thus, the APIM-peptide may have intrinsically high affinity for PCNA in the haematological cancer cells, while in primary non-proliferating monocytes from healthy blood donors external stress is needed to induce a similar response.

### Targeting PCNA in haematological cancer cells leads to changes in several canonical pathways including ubiquitin signaling and antigen presentation

We next explored if the metabolic signatures found in haematological cancer cells could be predicted from changes in cellular signaling. A refined variant of the MS-based multiple inhibitory bead (MIB)-assay was used for this purpose. This assay preferentially pulls down activated kinases via their increased affinity to kinase inhibitors (Duncan et al., 2012; Petrovic et al., 2017), and is a non-biased non-targeted approach for monitoring of the global “signalome”. When comparing the changes after APIM-peptide treatment in the haematological cancer cell lines JJN3, MC/CAR and NB4, we found a significantly increased pull down of 67 proteins and a significantly decreased pull down of 33 proteins common to all three cell lines (Figure 4A). INGENUITY pathways analysis (IPA) did not return any significantly enriched pathways from the list of increased proteins; however, the decreased proteins were found to be involved in ubiquitination, antigen presentations, calcium signaling and transport, and cellular responses to unfolded protein (Figure 4A).

**Figure 4.**
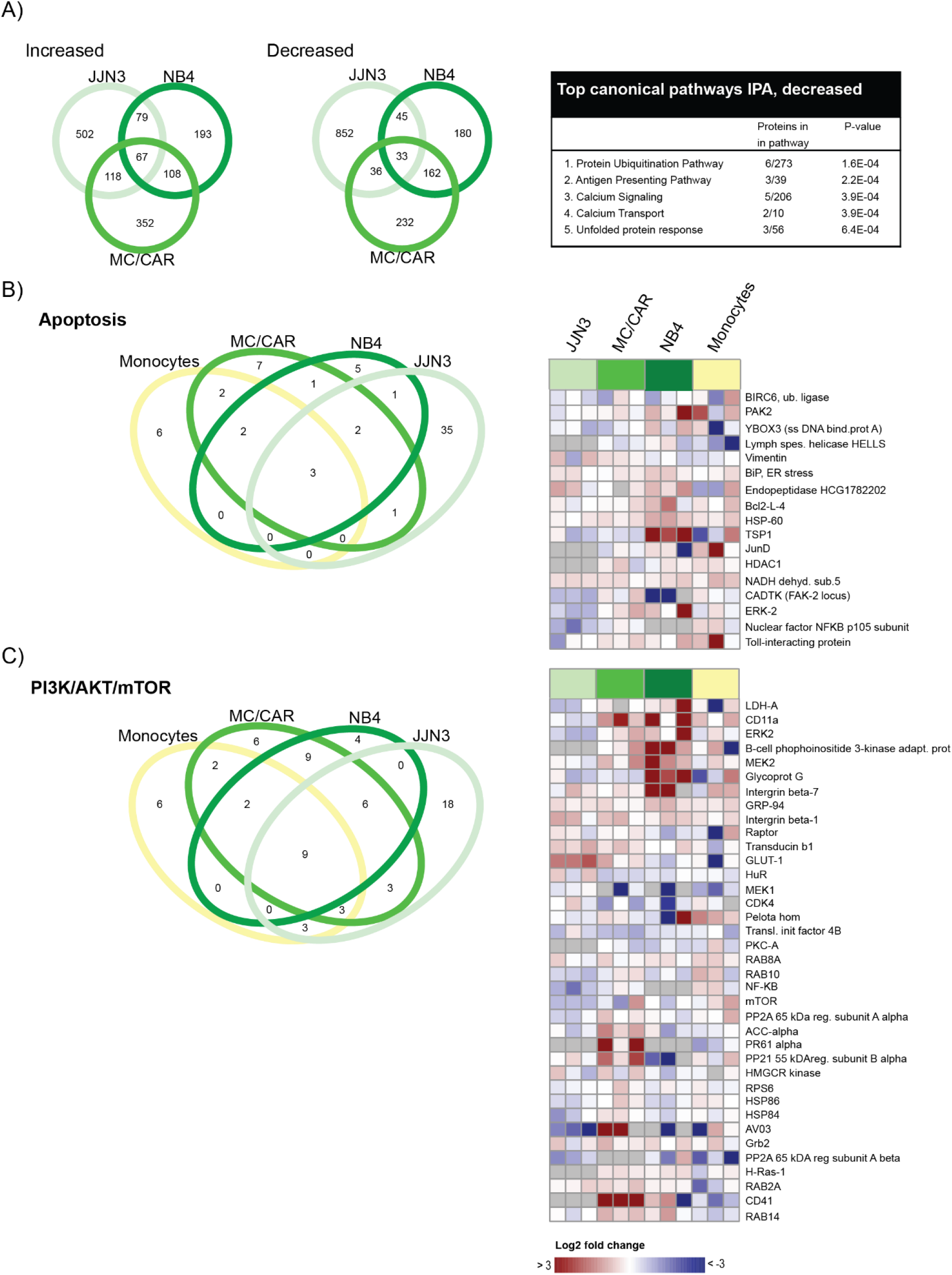
The signalome is strongly affected by targeting PCNA. Results of the MIB-assay of extract from haematological cells treated with APIM-peptide (JJN3: 6µM, MC/CAR, NB4, primary monocytes: 8 µM) for 4h. Data from three repeated experiments is shown, each presented as log_2_ fold change relative to untreated control. (**A**) Venn diagrams displaying number of proteins significantly changed in pull downs from treated cells relative to untreated control in JJN3, MC/CAR and NB4 according to the Wilcoxon Sign Rank test. Increased (left panel), decreased (mid panel) and results of IPA analysis of decreased proteins (right panel). (**B**) and (**C**) Left panel: Venn diagrams displaying number of proteins involved in apoptosis regulation and in the PI3K/AKT/mTOR pathway in treated cells relative to untreated control in JJN3, MC/CAR and NB4, and primary monocytes from three donors, according to the Wilcoxon Sign Rank test. Right panels: Heatmaps displaying proteins in the selected pathways that are significant changed from untreated control according to the Wilcoxon Sign Rank test in at least two cell types.

Based on previous results which have implicated both the PI3K/AKT/mTOR pathway and regulation of apoptosis in the response of haematological cells to APIM-peptide treatment (Müller et al., 2013; Olaisen et al., 2015), we next focused on changes in these pathways. Venn diagrams showing all proteins significantly changed in apoptosis and PI3K/AKT/mTOR pathways in JJN3, MC/CAR, NB4 and primary monocytes, and heat maps showing proteins which were significantly changed in at least two cell types (Figure 4B and C, left and right panel, respectively), indicated that the response of the two MM cell lines JJN3 and MC/CAR, were most similar. The APIM-peptide also influenced proteins in the MAPK, STAT and AMPK pathways in a similar manner in the two MM cells (Supplementary Figure S2) (full list of proteins in PRIDE, PDX017474). These results confirm that regulation of multiple central cellular signaling pathways are affected when the scaffold functions of PCNA is impaired (Olaisen et al., 2018; Olaisen et al., 2015; Sogaard, Blindheim, et al., 2018; Sogaard et al., 2019), and importantly, it suggests that MM cells share response patterns. Interestingly, the differences in responses after targeting PCNA in haematological cancer cells versus the primary monocytes were less pronounced at the signalome level compared to the metabolome level.

### Integrated analyses of metabolome, proteome, and transcriptome profiling support an important role of PCNA in regulation of metabolism in MM cells

We next explored the metabolic response to impairment of PCNA’s scaffold functions in more detail in the JJN3 MM cells compared to DU145 and Hek293 cells. Because APIM-peptide treatment enhanced the cytotoxic effects of cisplatin in cell lines sensitive to the APIM-peptide as a single agent (Supplementary Figure S1), we wanted to examine if this was reflected on the metabolic level. Most central carbon metabolite levels in JJN3 were strongly reduced already 4h after treatment with the APIM-peptide, an exception being the cysteine level, which increased 4-fold (Figure 5A). Combination treatment with cisplatin caused a similar response as the APIM-peptide alone, while cisplatin as a single-agent caused an overall upregulation of the metabolite pools. Thus, the changes in the metabolism is not a direct result of increased cytotoxicity. The same trends were observed after 8 and 24h in JJN3 (Supplementary Table S1D). However, similarly treated prostate cancer cells (DU145), which are sensitive to the APIM-peptide, and kidney cells (Hek293), which are not sensitive to APIM-peptide, showed minor changes in their central carbon metabolite pools even in presence of cytotoxic levels of cisplatin after 4, 8 and 24h (Supplementary Table S1E-F and Figure S1). This further suggests that the reduction of central metabolite pools upon APIM-peptide treatment is not directly linked to sensitivity to the peptide or cellular stress levels but is a haematological trait.

**Figure 5.**
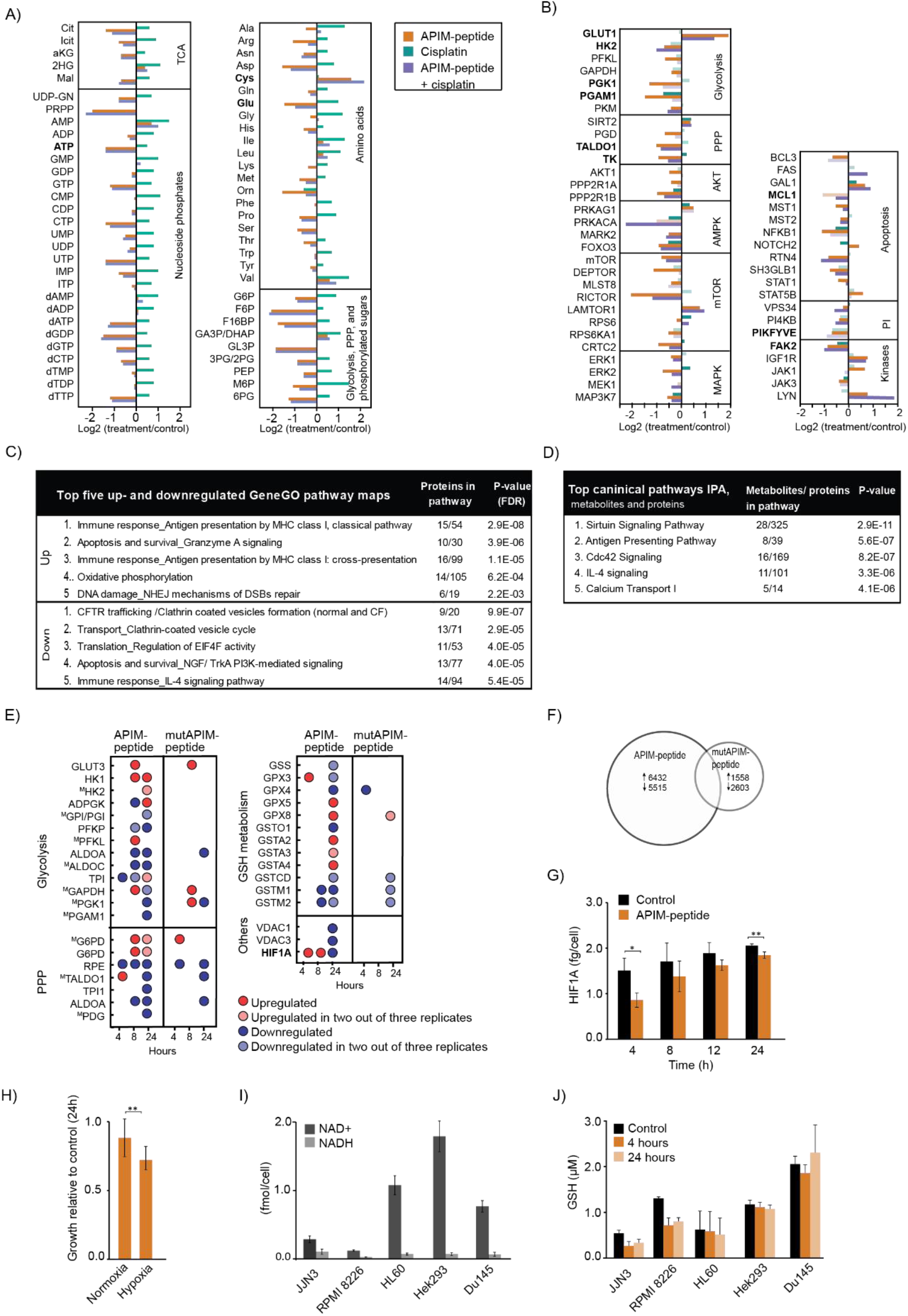
Metabolome, proteome and transcriptome profiling supports non-canonical role of PCNA in regulation of cellular signaling in JJN3 cells. (**A**) Log_2_ fold change of intracellular metabolite pools in JJN3 cells treated with APIM-peptide (8 µM), cisplatin (1 µM), or the combination given relative to control after 4h. The average of n = 4 from one representative out of three repeated experiments is shown. For abbreviations, see Supplementary Table S1A. (**B**) Results of MIB-assay of extract from JJN3 cells treated with APIM-peptide (6 µM), cisplatin (1 µM), or combination for 4h. Proteins in bold are mentioned specifically in the text. Data shown is mean of three repeated experiments, presented as log_2_ fold change relative to untreated control. Dark color bars indicate significant change from untreated control. (**C**) Top five up- and down-regulated GeneGO pathway maps based on significant changed proteins in the MIB-assays according to the Wilcoxon Sign Rank test (**D**) Top five pathways from IPA analysis of combined MIB and metabolite data sets (**E**) DE genes after APIM-peptide, or mutAPIM-peptide (6 µM) treatments. ^M^ denotes that the gene product also was detected by the MIB-assay. Average of three repeated experiments for 24h, and two repeated experiments for 4 and 8h (included only if same trend in both) is shown. (**F**) Number of DE genes in JJN3 cells after 24h of APIM- or mutAPIM-peptide (6 µM) treatment, (**G**) Average HIF1A protein levels (fg/cell) in JJN3 cells 4, 8, 12 and 24h after treatment with APIM-peptide (8 µM) ± SD, n = 3. (**H**) Average viability as measured by the MTT assay in JJN3 cells treated with APIM-peptide (6 µM) under atmospheric O_2_ tension (normoxia) or 1% O_2_ (hypoxia) at 24h ± SD, n = 10. Data from one representative out of two repeated experiments is shown, (**I**) Average basal NAD^+^ and NADH levels (fmol/cell) in JJN3, Hek293, DU145, RPMI 8226, and HL60 cells ± SD, n ≥ 4. (**J**) Average GSH levels after APIM-peptide (6 µM: JJN3, RPMI 8226, HL60, and DU145 or 8 µM: Hek293) treatment for 4 and 24h. Data based on triplicates from two (JJN3, Hek293, and RPMI 8226) or three (DU145 and HL60 repeated experiments. Details on replicate numbers are listed in Supplementary Table S2.

The effects of targeting PCNA with the APIM-peptide were evident even beyond central carbon metabolism. Four non-targeted LC-high mass accuracy MS methods covering both hydrophilic, hydrophobic, and charged metabolites were applied to profile treated JJN3 cells. Notably, and oppositely to the central metabolite pools which were generally down-regulated upon APIM-peptide treatment, most affected metabolites covered by non-targeted profiling were upregulated (Supplementary Figure S3A). Though the effects of cisplatin-APIM-peptide treatment were more pronounced than the effect of the APIM-peptide alone, the results of the two treatments were overlapping to a large extent (Supplementary Figure S3B).

To measure potential changes in the relative activity of central metabolic pathways, we next cultured APIM-peptide treated JJN3 cells in media with three differently labelled ^13^C-glucose isotopes. The labelling patterns obtained from two of the isotopes, 1^13^C_1_- and 1,2^13^C_2_-glucose yields information about glycolytic vs. PPP activities, while the third; fully labelled glucose, yields information about glucose consumption vs. consumption of other carbon sources, such as glutamine (Jang, Chen, & Rabinowitz, 2018). The ^13^C-isotope label enrichment from 1^13^C_1_- or 1,2^13^C_2_-glucose revealed no differences between treatments. However, the labelling from fully labelled glucose in metabolites of lower glycolysis, i.e. 2-/3PG and in citrate were slightly lower in APIM-peptide treated cells already after 4h. This also applied to the PPP intermediate S7P (Supplementary Table S1G), supporting that the intracellular flux distribution of central metabolic pathways is affected by the drop in glucose uptake in these cells (Figure 2A).

In support of perturbed glycolysis, we observed from the MIB-assay that pull-down of glycolytic and PPP proteins such as hexokinase 2 (HK2), phosphoglycerate kinase 1 (PGK1), phosphoglycerate mutase 1 (PGAM1), transaldolase (TALDO1), and transketolase (TK), were strongly reduced compared to untreated control in extracts from APIM-peptide or combination-treated JJN3 cells (Figure 5B, bolded proteins). Cisplatin as a single agent had a smaller effect than the APIM-peptide and the combination, and as observed on the metabolome level, the combination treatment was more similar to APIM-peptide alone than to cisplatin alone. This differs from solid tissue cancer cells: in both an APIM-peptide sensitive (Um-Uc-3) and a less/not sensitive (T24) bladder cancer cell, the effect of the APIM-peptide as a single agent was limited, while the peptide enhanced the effects of cisplatin in both cell lines at the metabolome, signalome and the transcriptome level (Sogaard et al., 2019).

A reduction in the levels of several proteins important for regulation of metabolism, e.g. proteins in the AKT, AMPK, mTOR and MAPK pathways and several anti-apoptotic proteins, were detected upon APIM-peptide treatment of JJN3 cells (Figure 5B). One example of the latter is MCL1, shown to be vital for survival of JJN3 cells (Gong et al., 2016). Out of the other signaling proteins, the most evident changes were the reduction of PI kinases, including PIKFYVE, important for vesicular transport, and the non-receptor tyrosine kinases FAK2, known to activate MAPK signaling and a potential drug target in MM (Meads et al., 2016). Additionally, increased pull-down of important endoplasmic reticulum (ER) proteins/antioxidants, e.g. PRDX4 and ERO1A which are important for redox balance in ER, and proteins involved in DNA repair, e.g. the DNA strand break sensors PARP1 and XRCC6/Ku70, were detected in APIM-peptide treated cells (Supplementary Figure S4A, KEGG pathway analysis for protein processing in ER is shown in Supplementary Figure S5A). Taken together, the changes detected on the level of the signalome show that targeting the scaffold function of PCNA affects several signaling networks important for regulation of metabolism, redox balance, oxidative protein folding and apoptosis.

GeneGo pathway analysis of changed proteins, and IPA analysis combining changed proteins and metabolites, pointed to alterations in pathways regulating oxidative phosphorylation, immune response/antigen presentation, and vesicle and calcium transport in APIM-treated JJN3 (Figure 5C and D). Calcium signaling and antigen presentation were also two of the top five affected canonical pathways revealed by IPA analysis of proteins changed in JJN3, MC/CAR and NB4 cells (Figure 4A). Pull down of most of the proteins involved in oxidative phosphorylation were increased upon APIM-peptide treatment of JJN3 cells (Supplementary Figure S4), and many of these were membrane proteins. PI kinases are important for both cellular signaling and vesicular transport, and several of these, including PIKFYVE, are downregulated in our dataset (Figure 5B). Impaired vesicle transport could therefore lead to an increase in pull-down of membrane proteins due to increased concentrations of these proteins in the cell extracts. Accordingly, when comparing our dataset from the MIB-assay to a list of membrane proteins, the “membranome” (Uva et al., 2010), a clear APIM-peptide signature was detected and ∼80% of these proteins were upregulated (Supplementary Figure S4B and C). APIM-peptide-mediated downregulation of vesicular trafficking in JJN3 is also illustrated by KEGG pathway analysis (Supplementary Figure S5B).

Transcriptome profiling was performed to complement the metabolome and proteome data sets. The response of JJN3 cells treated with APIM-peptide compared to mutAPIM-peptide verified that most of the differentially expressed (DE) genes were linked to the ability of the APIM-peptide to interact with PCNA. For example, APIM-peptide treatment led to a higher number of DE genes in total, as well as an increasing number of DE genes involved in glycolysis, PPP and GSH metabolism over time, and this was not, or too a much lower extent, detected in mutAPIM-peptide treated cells (Figure 5F and E, respectively). Whereas large changes in the metabolome and proteome were evident after 4h, the effect on gene expression was more prominent after 24h. This suggest that the early effects observed after targeting PCNA are more direct consequences of inhibiting protein - PCNA interactions, and that the effects observed on the transcriptome are indirect long-term effects of changes in signaling.

### Targeting PCNA impairs the HIF1A response and the redox balance in multiple myeloma cells

The transcription levels of HIF1A, a transcription factor known to upregulate glycolysis and to be a mediator of cells response to hypoxic conditions, was initially increased (4 and 8h), but reduced 24h after APIM-peptide treatment (Figure 5E). Protein levels of HIF1A are known to be rapidly regulated via post-translational modifications (Salceda & Caro, 1997). When protein levels of HIF1A were measured, reduced protein levels of HIF1A were found already after 4h (Figure 5G). Further supporting the ability of the APIM-peptide to perturb a normal HIF1A response, the treatment had a stronger effect on JJN3 cell growth under hypoxic than normoxic conditions (Figure 5H).

IPA analysis of the combined protein and metabolite data sets from JJN3 cells identified the NAD^+^ -dependent de-acetylase Sirtuin signaling pathway as the most significantly modified. Interestingly, the basal levels of NAD^+^ were at least three times lower in the MM cell lines JJN3 and RPMI 8226 than in the other cell lines examined (Figure 5I). Targeting the scaffold functions of PCNA with the APIM-peptide reduced the NAD-levels in all cell lines tested. These changes were attributed to APIM - PCNA interactions as the mutated APIM-peptide and the peptide containing only the cell penetrating part of the APIM-peptide caused a smaller, or close to no effect, respectively (Supplementary Figure S6). NAD^+^/NADH and glutathione (GSH)/oxidized GSH (GSSH) are redox couples important in the cellular defense against oxidative stress. Thus, lower basal NAD^+^ levels in MM cells could be caused by a high redox demand due to high synthesis of immunoglobulins. Interestingly, GSH levels were reduced in the two MM cell lines upon APIM-peptide treatment, while not in the other cell lines tested (Figure 5J). Biosynthesis of GSH requires cysteine, ATP and glutamate. Levels of glutamate and ATP are strongly reduced upon APIM-peptide treatment in JJN3 cells, while the cysteine level is increased (Figure 5A). This may indicate impaired GSH synthesis due to low ATP levels upon APIM-treatment in MM cells. Interestingly, GSH synthetase (GSH-S) contains an APIM sequence (Olaisen et al., 2018). Taken together, these results support impaired cellular defense against oxidative stress in the MM cells and indicate that the intrinsic low levels of NAD^+^ may be a key point for explaining the hypersensitivity to APIM-peptides in MM cells.

## Discussion

While nutrient uptake normally is strictly regulated, a metabolic phenotype characterized by high rate of glycolysis even in the presence of oxygen, i.e. “the Warburg effect”, is frequently found in cancers. This reprogramming/switch of energy metabolism is thought to support the increased need for energy, biosynthetic precursors, and reducing power required for rapid proliferation. Metabolic reprogramming is now firmly established as a hallmark of cancer (Hanahan & Weinberg, 2011). Several attempts have been made to attack this malignant transformation; however, no inhibitors of glycolysis are yet approved as anticancer agents. In this study, we have examined effects of inhibiting PCNA - protein interactions in cells of different origin, and we show that PCNA’s scaffold role is more prominent for regulation of metabolism in haematological cells than in cells from solid tissue. Figure 6 summarizes the observed changes in levels of central carbon metabolites and selected enzymes and regulatory proteins in the MM cell line JJN3. Overall, the reduction in both metabolites and proteins important for glucose metabolism is striking. Reduced levels of HK2, PGAM1, and PGK1 were detected after APIM-peptide treatment, all of which are glycolytic proteins regulated by, or regulating the PI3K/AKT/mTOR pathway and shown to be important for tumor growth and metastasis (Gao & Han, 2018; Liu et al., 2018; Yu et al., 2017). The AMPK pathway is normally activated by a lowered ATP/AMP ratio and serve to increase glucose uptake in order to maintain cellular ATP levels (Faubert, Vincent, Poffenberger, & Jones, 2015). However, reduced activation of the AMPK pathway upon impairment of the scaffold functions of PCNA, may impair the normal regulation so that cells become depleted of ATP. One key metabolic link for the MM cell hypersensitivity to the APIM-peptide treatment, seems to be low intrinsic concentration of NAD^+^, i.e. low reducing power. Notably, analyzing the global metabolite pool, we found that most metabolite levels increased after APIM-peptide treatment, which strongly indicates that the reduction of central metabolite levels is caused by inhibition of the scaffold function of PCNA, and not a general shut-down of the cells.

**Figure 6.**
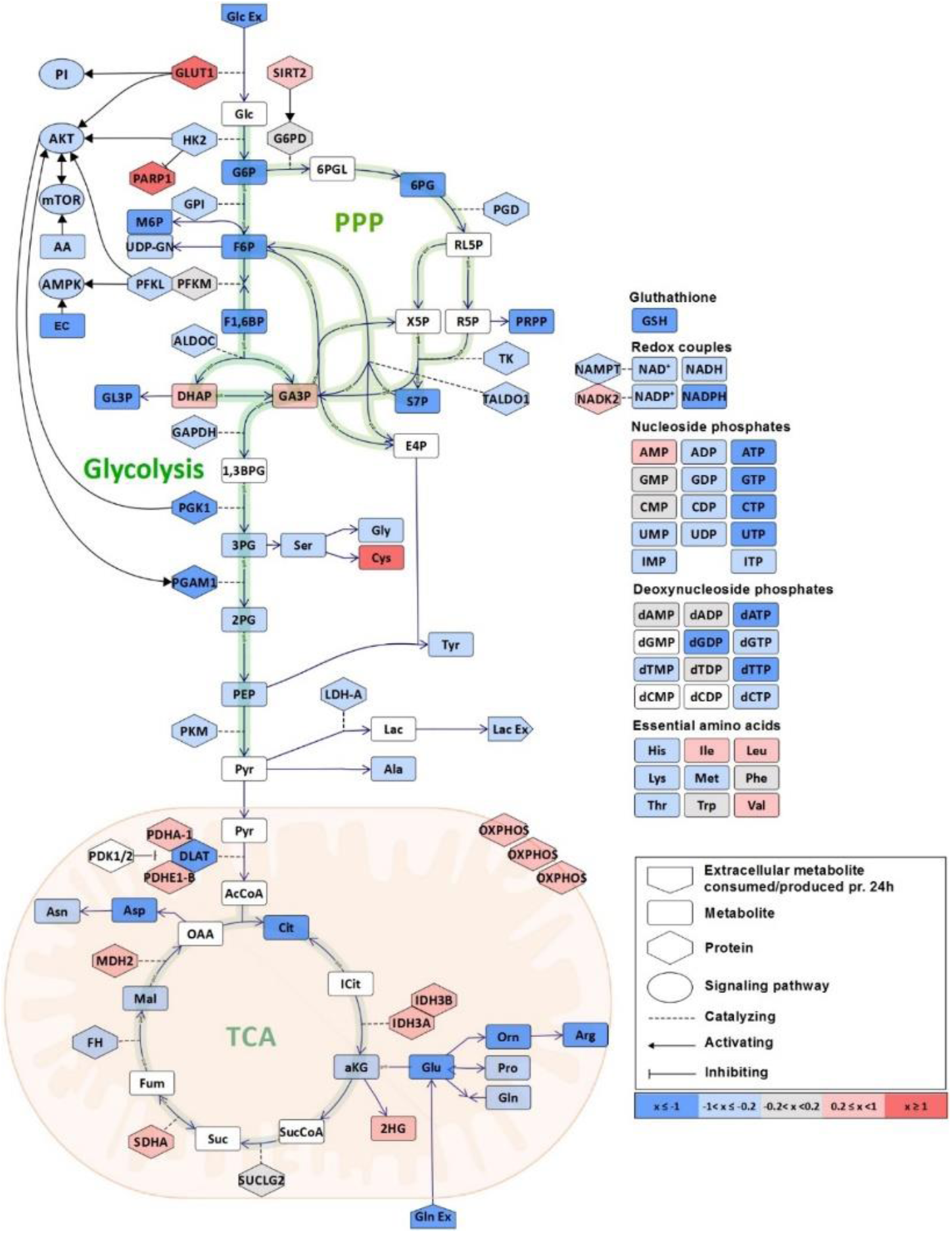
Targeting PCNA with APIM-peptide alters metabolite pools, metabolic enzymes and associated signalling pathways in JJN3 cells. Simplified schematic overview of central carbon metabolism showing log_2_ fold change of levels of glycolytic, TCA and PPP intermediates, phosphorylated sugars, amino acids and (deoxy)nucleoside phosphates with associated proteins and pathways in JJN3 cells treated with APIM-peptide (6/8 µM) relative to control after 4h. White color indicates that metabolite/protein was not measured or detected. For grouped metabolites and pathways, the average log_2_ fold change is presented. The figure is based on data presented in Figures 2-4. For abbreviations, see Supplementary Table S1A.

A major problem when studying cytosolic roles of PCNA is to distinguish these effects from its canonical roles in DNA replication and repair. We have shown that APIM-peptide as a single agent leads to apoptotic responses, but it had no effect on cell cycle distribution in JJN3 (Müller et al., 2013) and DU145 cells (Sogaard, Moestue, et al., 2018). Furthermore, we recently showed that targeting PCNA with the APIM-peptide in combination with receptor tyrosine kinase inhibitors increased their anti-cancer activities in breast, colon and bladder cancer cells, and changed their cellular signaling responses (Sogaard et al., 2019). These results support a cytosolic role of PCNA as no DNA damage/replication stress was induced. Further, integration of data presented herein and in another recent study (Sogaard, Blindheim, et al., 2018) indicates that prostate (DU145), kidney (Hek293), and bladder (Umuc-3, T24) cells are less dependent on a PCNA-governed regulation of pathways controlling glycolysis than haematological cells, and/or that they have an increased ability to reprogram signaling networks to avoid these changes. However, even though levels of glycolytic intermediates and nucleoside phosphates were not/or less affected by APIM-peptide treatment in DU145, Hek293 and T24 cells (Figure 2A), their clear reduction in NAD^+^ and NADH pools upon APIM-peptide treatment suggest that PCNA is important for the regulation of cellular metabolism also in these cells (Supplementary Figure S6). We also detect clear effects on bladder cancer metabolism by targeting PCNA with the APIM-peptide (Sogaard, Blindheim, et al., 2018); however, these effects differ from those detected in haematological cells. Tissue- or cell type specific roles of PCNA and a link to glucose metabolism has also been suggested previously (Naryzhny & Lee, 2010; Ohayon et al., 2016).

Multiple kinases (∼90) contain a potential PCNA-interacting motif (Olaisen et al., 2018); thus, the effects of targeting PCNA measured on the level of metabolism could be an indirect consequence of reduced activity of multiple signaling pathways (e.g. PI3K, AKT, mTOR, AMPK, MAPK). However, it could also be a direct consequence of inhibition of interactions between PCNA and APIM and PIP-box containing proteins directly participating in glycolysis: i) Glucose-6-phophate dehydrogenase (G6PD), the first and rate-limiting step in the PPP contains a PIP-box. ii) Pyruvate dehydrogenase kinase (PDHK1) also contains a PIP-box and was recently implicated in the regulation of glycolysis in T-cells (Menk et al., 2018). PDHK1 is rapidly activated by T-cell receptors and involved in turning glucose consumption towards aerobic glycolysis by phosphorylation and inactivation of PDH and prevention of mitochondrial import of pyruvate. iii) Enolase-1 (ENO1), catalyzing the enzymatic conversion of 2-phosphoglycerate (2PG) to phosphoenolpyruvate (PEP), was recently shown to be regulated via the AMPK-AKT pathway and also, ENO1 contain APIM (Dai et al., 2018; Olaisen et al., 2018). Inhibition of an interaction with the scaffold protein PCNA for either of the above-mentioned enzymes might have a direct effect on central carbon metabolism and could represent an additional level of metabolic regulation. Most likely, the effects observed on metabolism by APIM-peptide treatment are a combination of both direct inhibition of interactions between glycolytic enzymes and PCNA, and indirect effects of altered cellular signaling. One way to explore this further is by site specific mutation of PCNA interacting motifs (e.g. by CRISPR/Cas9). However, due to the multi-level regulation of central metabolism it might be that single mutants, e.g. of G6PD, PDHK1, ENO1, do not show a phenotype neither at local nor global metabolite levels. Thus, series of mutations might be needed for appearance of a distinct phenotype.

The APIM-peptide interacts with the same region on PCNA as PIP-containing peptides (Müller et al., 2013; Sebesta et al., 2017). The *in vitro* APIM-peptide – PCNA interaction is much weaker than the canonical PIP-box peptide derived from p21; however, the current hypothesis is that APIM-peptide affinity is increased under cellular stress (Ciccia et al., 2012; Gilljam et al., 2009; Olaisen et al., 2015; Prestel et al., 2019). Previously, we measured a strong inhibition of cytokine secretion from LPS stimulated monocytes after treatment with the APIM-peptide at doses that did not induce apoptosis (Olaisen et al., 2015). Here we show that the APIM-peptide treatment of LPS activated monocytes lead to a “shutdown” of the central metabolism; however, this effect was not detected in absence of LPS stimulation. These results are in line with the hypothesis that cellular stress increased the affinity of PCNA to APIM, thereby favoring the APIM-proteins over PIP-box containing proteins for interaction. Thus, stress is mediating a switch in PCNA’s proteins partners which is important for the cellular stress response. Modulation of the immune reaction only in “activated” immune cells could be advantageous in the tumor microenvironment where pro-inflammatory cytokines might “fuel” the tumors (Farajzadeh Valilou, Keshavarz-Fathi, Silvestris, Argentiero, & Rezaei, 2018). Such modulation of the immune reactions could possible explain the disease stabilization observed in certain patients in the current ongoing phase I study of the lead APIM-peptide ATX-101.

We are only beginning to gain knowledge about the non-canonical roles of PCNA, but recent results, including this study, have established that the cytosolic roles of PCNA are complex and vary in different cell types (Müller et al., 2013; Naryzhny & Lee, 2010; Ohayon et al., 2016; Olaisen et al., 2015; Rosental et al., 2011; Sogaard, Blindheim, et al., 2018; Sogaard, Moestue, et al., 2018; Sogaard et al., 2019; Witko-Sarsat et al., 2010; Yin et al., 2015). Experimental studies of multifunctional proteins are extremely challenging, and robust quantitative methodologies for measurements at the metabolome and protein level are needed. The integrated analyses of viability, metabolome, proteome and transcriptome profiling assays presented herein strongly indicates that protein - PCNA interactions have special roles in regulation of central carbon metabolism in haematological cells.

## Material and Methods

### Methods details

#### Peptides and stressors

APIM-peptide (Ac-MDRWLVKWKKKRKIRRRRRRRRRRR), mutAPIM-peptide (Ac-MDRALVKWKKKRKIRRRRRRRRRRR) (Müller et al., 2013), and R11 (Ac-RRRRRRRRRRR) (Olaisen et al., 2015) (Innovagen, Sweden) were used. Monocytes were stimulated with Lipopolysaccharide (LPS) (Sigma-Aldrich) for 4h, and cell lines were treated with cisplatin (Hospira).

#### Cell lines and assay conditions

Cell densities were optimized for each individual assay and corresponding sampling time points, and peptide doses were adjusted to cell density to obtain the same number of annexin-positive cells in all experiments. These data are listed in Supplementary Table 2, also listing number of independent cultures analyzed (n) and number of repeated experiments. JJN3 (Müller et al., 2013) and RPMI 8226 (ATCC CCL-155) (MM), HL60 (ATCC CCL-240) and NB4 (Kind gift from professor Stein Døskeland, University of Bergen, Norway) (AML), and DU145 (ATCC HBT-81, prostate cancer) cells were cultured in RPMI 1640 (Sigma-Aldrich), MC/CAR (ATTC® CRL-8083) (MM) were cultured in IMDM (Sigma-Aldrich), and Hek293 (ATCC CRL-1573, embryonic kidney) were cultured in DMEM high glucose (Sigma-Aldrich), all supplemented with 2mM glutamine (Biochrome), 100 µg/mL gentamicin (Sigma-Aldrich), 2.5 µg/mL amphotericin (Sigma-Aldrich), and fetal bovine serum (Sigma-Aldrich); 10% in DMEM high glucose and RPMI 1640, and 20% in IMDM. Cells were maintained at 37°C in a humidified atmosphere of 5% CO_2_.

#### Primary monocytes

Peripheral blood monocytes were isolated and cultured from three A +/- buffy coats (Blood Bank, St. Olav’s University Hospital, Norway) as described in (Olaisen et al., 2015).

#### Viability assay

The MTT assay was performed as described in (Gilljam et al., 2009). Cells were incubated in a humidified atmosphere of 5% CO_2_ and either atmospheric O_2_-levels (normoxia) or 1% O_2_ (hypoxia), the latter in a C-chamber incubator sub chamber controlled by a ProOx C21 compact O_2_ and CO_2_ sub chamber controller from BioSpherix.

#### Measurements of extracellular metabolites

Extracellular glucose, lactate and glutamine was measured as described in (Sogaard, Blindheim, et al., 2018).

#### Targeted mass spectrometric metabolite profiling

JJN3, RPMI 8226, MC/CAR, HL60 and NB4 cells were sampled and extracted as described in (Kvitvang & Bruheim, 2015); by fast filtering at a maximum vacuum pressure of 250 mbar below the ambient pressure including two consecutive rinsing steps with 10 ml saline and MQ-water, respectively. Two filters were pooled in 13 ml MQ-water:acetonitrille (1:1, v/v). Hek293 and DU145 cells and primary monocytes were sampled as described in (Kvitvang, Kristiansen, & Bruheim, 2014). All cell suspensions were immediately quenched in LN_2_, and extracted, lyophilized and cleared of protein as described in (Kvitvang & Bruheim, 2015), before reconstitution in compatible solvents for downstream analysis. TCA intermediates and phosphorylated metabolites were prepared for and quantified by capIC-MS/MS as described in (Kvitvang et al., 2014), with the modifications described in (Stafsnes, Rost, & Bruheim, 2018) employed for the haematological cells. Amino acids in DU145 and Hek293 extracts were quantified by GC-MS/MS as described in (Kvitvang, Andreassen, Adam, Villas-Boas, & Bruheim, 2011). Amino acids in two out of three replicates of JJN3 extracts were derivatized and analyzed as described in (Sogaard, Blindheim, et al., 2018). Amino acids in all remaining haematological cell extracts were derivatized and analyzed as described (Røst et al., 2020). Lactate and pyruvate in MC/CAR and NB4 were derivatized by a protocol adapted from (Tan, Lu, Dong, Zhao, & Kuo, 2014), and analyzed as described in (Sogaard, Blindheim, et al., 2018).

#### LC-MS/MS analysis of NAD pools

Intracellular NAD was extracted from pellets (4°C, 150 x g, 3 min) by shaking (2 min, 80 °C, 1500 rpm) in 300 µL water-acetonitrile with 10 mM acetic acid, adjusted to pH 9.0 with ammonium hydroxide (80°C, 10-90 v/v). Residual lipids and proteins were cleared from the extracts by spin filtration in 3-kDa-molecular-weight spin cut-off filter (VWR) added 3 mg C_18_-material (Waters). NAD levels were quantified applying a Waters ACQUITY I-Class UPLC/Xevo TQ-S triple quadrupole MS system operated in in negative electrospray mode. Samples (5 µl) were injected onto a Waters ACQUITY UPLC BEH-Amide 1.7 µm 2.1×100mm column, maintained at 40°C and eluted with mobile phases (A) water-acetonitrile (60-40 v/v), and (B) water-acetonitrile (10-90 v/v), both added 10 mM acetic acid and adjusted to pH 9.0 with ammonium hydroxide. The following gradient was applied with a flow rate of 0.4 ml/min: 0-0.5 min: 95% B, 0.5-2 min: 95-70% B, 2-6.5 min: 70-40% B, 6.5-7 min: 40-1% B, 7-8 min: 1% B, 11 min: end. Analytical grade standards (Sigma-Aldrich) were prepared fresh in mobile phase B, and both standards and extracts were analyzed within four hours.

#### Non-targeted mass spectrometric metabolite profiling

Extracts of JJN3 cells were prepared for analysis by high resolution high mass accuracy mass spectrometry as described for targeted mass spectrometric metabolic profiling. Three different chromatographic separation techniques were applied to maximize the coverage; capIC, reverse phase (RP) LC and hydrophobic interaction chromatography (HILIC), the two latter performed on a Waters ACQUITY I-Class UPLC. Chromatographic separation was interphased to a Waters SYNAPT HDMS QTOF mass spectrometer. **RP LC-MS:** Analysis of non-polar to medium polar metabolites was performed twice, employing positive and negative electrospray ionization. Samples (2 µl) were injected onto a Waters ACQUITY HSS T3 C18 1.7 µm 2.1*100 mm column, maintained at 40°C and eluted with mobile phases (A) water and (B) acetonitrile, both added 0.1% acetic acid. The following gradient was applied with a flow rate of 0.5 ml/min: 0-0.5 min: 1% B, 0.5-2 min: 1-15% B, 2-3.5 min: 15-50% B, 3.5-4.5 min: 50-99%B, 4.5-5 min: 95% B, 5-5.1 min: 95-1% B, 7 min: end.

#### HILIC-MS

Analysis of polar metabolites was performed employing negative electrospray ionization. Samples (2µl) were injected onto a Waters ACQUITY BEH amide 1.7 µm 2.1*100 mm column, maintained at 40°C and eluted with (A) water-acetonitrile (60-40, v/v%) and (B) water-acetonitrile (5-95, v/v%), both added 10 mM ammonium acetate and adjusted to pH 9.0 with ammonium hydroxide. The following gradient was applied with a flow rate of 0.4ml/min: 0-1 min: 99% B, 1-2 min: 90-70% B, 2-3.5 min: 70-55% B, 3.5-3.6 min: 55-1% B, 3.6-3.9 min: 1% B, 3.9-4.0 min: 1-99 % B, 6 min: end. **CapIC-MS:** Chromatographic separation of anionic metabolites was performed by capIC as described in (Kvitvang et al., 2014). The mass spectrometer was operated in negative electrospray mode.

#### ^13^C-labeling experiments

JJN3 cells were seeded in RPMI 1640 medium without glucose (Gibco) supplemented with either 1^13^C_1_-, 1,2^13^C_,2_- or U^13^C_6_-glucose (2 g/l, 99%, Cambridge Isotope Laboratories) and incubated over night before treatment. Sampling and extraction were performed as described for targeted mass spectrometric profiling, and analyzed with capIC separation as described, with a dedicated MS/MS method programmed for detection of all possible ^13^C-isotopomers in selected intermediates of glycolysis, the PPP, the TCA cycle and in all nucleoside triphosphates.

#### GSH levels

GSH levels were measured using the GSH-Glo™ Glutathione Assay (Promega) according to the manufacturer’s instructions.

#### HIF1A levels

Whole protein extracts were prepared from JJN3 pellets (4°C, 150 x g, 5 min) according to the cell extraction protocol for ELISA sample preparation by ThermoFisher Scientific. HIF1A levels were measured in duplicates applying a HIF1A Human ELISA kit (Invitrogen) according to the manufacturer’s instructions. Measured HIF1A levels were normalized to cell density at the time of sampling

#### Multiplexed inhibitor bead (MIB)-assay

Cell extracts for the MIB-assay (kinase enrichment) were prepared as described (Petrovic et al., 2017).

#### Whole genome gene expression analysis

Total RNA was isolated from snap-frozen pellets employing the RNeasy kit (Qiagen). Genome-wide gene expression profiling analysis was performed as described (Sogaard, Moestue, et al., 2018).

### Quantification and statistical analysis

#### Statistical analysis

Statistical tests are given in the relevant figure legends. These includes ANOVA, post hoc Turkey’s test, student’s T-test, and Wilcoxon Sign Rank test.

#### Gene expression analysis

Raw data was normalized and analyzed using GeneSpring 12.6 – GX. Probes were filtered by Flags, and fold change >=1.1. Similar up and downregulated genes from duplicate/triplicate experiments were extracted.

#### MIB-assay data analysis

Proteins were quantified by processing MS data using Max Quant v 1.6.4.0. (Tyanova, Temu, & Cox, 2016). Preview 2.3.5 (Falkner, Falkner, Yocum, & Andrews, 2008) was used to inspect the raw data to determine optimal search criteria. The following search parameters were used: enzyme specified as trypsin with maximum two missed cleavages allowed; acetylation of protein N-terminal, oxidation of methionine, deamidation of asparagine/glutamine and phosphorylation of serine/threonine/tyrosine as dynamic post-translational modification. These were imported in Max Quant which uses m/z and retention time (RT) values to align each run against each other sample with a minute window match-between-run function and 20 mins overall sliding window using a clustering-based technique. These were further queried against the Human proteome including isoforms downloaded from Uniprot (www.uniprot.org/proteomes/UP000005640) in October 2018 and Max Quant’s internal contaminants database using Andromeda built into Max Quant. Both Protein and peptide identifications false discovery rate (FDR) was set to 1%, thus only peptides with high confidence were used for final protein group identification. Peak abundances were extracted by integrating the area under the peak curve. Each protein group abundance was normalized by the total abundance of all identified peptides for each run and protein by calculated median summing all unique and razor peptide-ion abundances for each protein using label-free quantification (LFQ) algorithm (Cox et al., 2014) with minimum peptides ≥ 1. LFQ values for all samples were combined and log-transformed with base 2 and the transformed control values were subtracted. The resulting values reflecting the change relative to control for each condition were subjected to two-sided non-parametric Wilcoxon Sign Rank Test (Gibbons & Chakraborti, 2010) as implemented in MATLAB R2015a (Math Works Inc) in order to check the consistency in directionality of the change, namely a negative sign reflecting decreased and positive sign reflecting increased expression of respective protein group. The choice of this non-parametric test avoids the assumption of a certain type of null distribution as in Student’s t-test by working over the Rank of the observation instead of observation value itself. Further, it also makes it robust to outliers and extreme variations noticed in observed values. DE protein groups were identified at p 0.25. The Uniprot accession IDs of these DE were mapped to pathways (www.wikipathways.org/index.php/Download Pathways version wikipathways-20191010-gmt-Homo_sapiens.gmt) using R (https://www.R-project.org/) libraries, org.Hs.eg.db and clusterProfiler (www.liebertpub.com/doi/10.1089/omi.2011.0118). Venn diagrams were built using the R package limma (Ritchie et al., 2015). Online Ingenuity® Pathway Analysis™ software (QIAGEN Inc., www.qiagenbioinformatics.com/products/ingenuitypathway-analysis) was used to combine with metabolomics data for annotation, visualization and integrated discovery of canonical pathways and other functional analysis.

### Metabolite analysis

#### Quantification of extracellular metabolites

Extracellular glucose, lactate and glutamine was quantified as described in (Sogaard, Blindheim, et al., 2018).

#### Quantification of intracellular metabolites

Downstream data processing was performed in TargetLynx application manager of MassLynx 4.1 (Waters). Ion abundances were normalized to total ion abundance in DU145 and Hek293 extracts. Metabolite levels in JJN3, RPMI 8226, MC/CAR, HL60, NB4, and primary monocyte extracts were absolutely quantified by interpolation from calibration curves prepared by serial dilutions of standards calculated by least squares regression. Extract concentrations were normalized to cell density at the time of sampling.

#### ^13^C labelling experiments

Ion abundances of all possible isotopologues were quantified in TargetLynx application manager for MassLynx 4.1 to allow for calculation of relative summed fractional labelling (SFL).

#### Non-targeted mass spectrometric metabolite profiling

Raw data files were processed with Progenesis QI software (Waters), with the threshold for peak picking set to the 500 to 1000 most abundant masses. The datasets were trimmed of ions which abundance were highest in blank samples, or which abundance was > 10% of ‘pooled quality control’ abundance in blank samples. Finally, ion abundances were normalized to cell density at the time of sampling.

## Data and code availability

The microarray experiments are MIAME (Minimum Information About a Microarray Experiment) compliant and have been deposited in the ArrayExpress database (http://www.ebi.ac.uk/arrayexpress/) under accession number E-MTAB-5644.

The MS proteomics data has been deposited to the ProteomeXchange Consortium via the PRIDE (Vizcaino et al., 2016) partner repository with the dataset identifier, project ID PXD011044; raw data JJN3 and project ID PDX017474; raw data the rest of the cell lines and monocytes, all results and program codes.

## Acknowledgements

We thank Anala Nepal, Siri Bachke and Ida Eide Langørgen for technical assistance. Proteomics were performed at Proteomics and Modomics Experimental Core Facility (PROMEC), NTNU, and NorStore/Notur Project NS9036K was used for data storage and computation services. The MS based metabolite profiling and NMR-analysis of extracellular metabolites were performed at the NTNU NV-faculty MS and NMR facilities, respectively.

## Author contributions

L.M.R., C.O., P.B., and M.O. designed the experiments, L.M.R., C.O., A.S., H.F.K., V.P., P.B., and M.O. developed the methodology, L.M.R., C.O., A.N., V.P., H.F.K. and P.B. performed acquisition of data, L.M.R., C.O., A.S. A.N., V.P., P.B., and M.O. performed analysis and interpretation of the data and L.M.R., C.O., P.B. and M.O wrote the paper.

## Abbreviations

AML: acute myeloid leukemia;
APIM: AlkB homologue 2 PCNA-interacting motif;
APIM-peptide: APIM-containing peptide;
mutAPIM-peptide: mutated APIM-peptide;
capIC: capillary ion chromatography;
DE: differentially expressed;
LC: liquid chromatography;
LPS: Lipopolysaccharide;
IPA: INGENUITY pathways analysis;
MIB: multiplexed inhibitor beads;
MM: multiple myeloma;
MS: mass spectrometry;
MS/MS: tandem mass spectrometry;
PCNA: proliferating cell nuclear antigen;
PIP-box: PCNA-interacting protein-box;
PI: phosphatidylinositol;
PPP: pentose phosphate pathway;
PTM: posttranslational modifications;
ROS: reactive oxygen species;
TCA: tricarboxylic acid.

