## Supplementary Figures and Tables for "PCNA has specific functions in regulation of metabolism in haematological cells"

#### Supplementary Figure S1, related to Figure 2 and 5

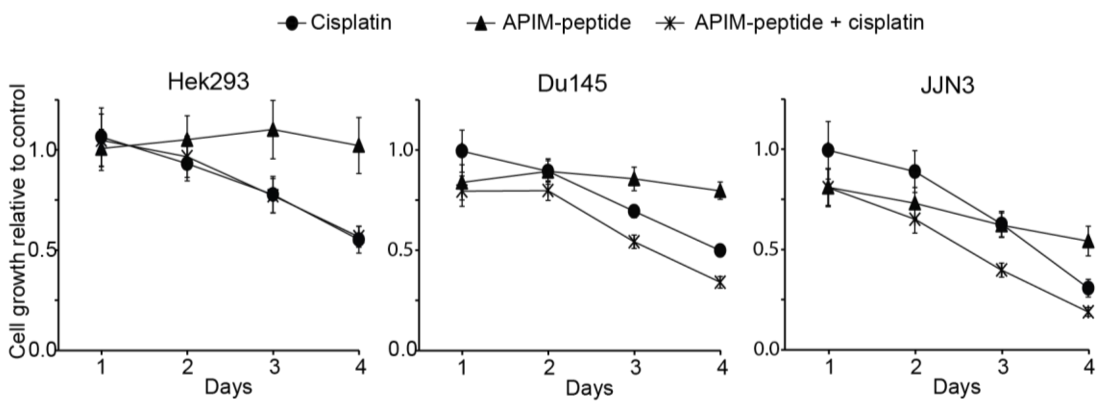


**Supplementary Figure S1. APIM-peptide sensitivity differs between cell lines.** Viability assay (MTT) of Hek293, DU145, and JJN3 cell lines treated with APIM-peptide (8 µM for Hek293; 6 µM for DU145 and JJN3), cisplatin (1 µM), or the respective combination relative to untreated control. Cells were harvested on day 1-4 after treatment. The average of two repeated experiments, each with n ≥ 6 is shown.

### Supplementary Figure S2, related to Figure 4

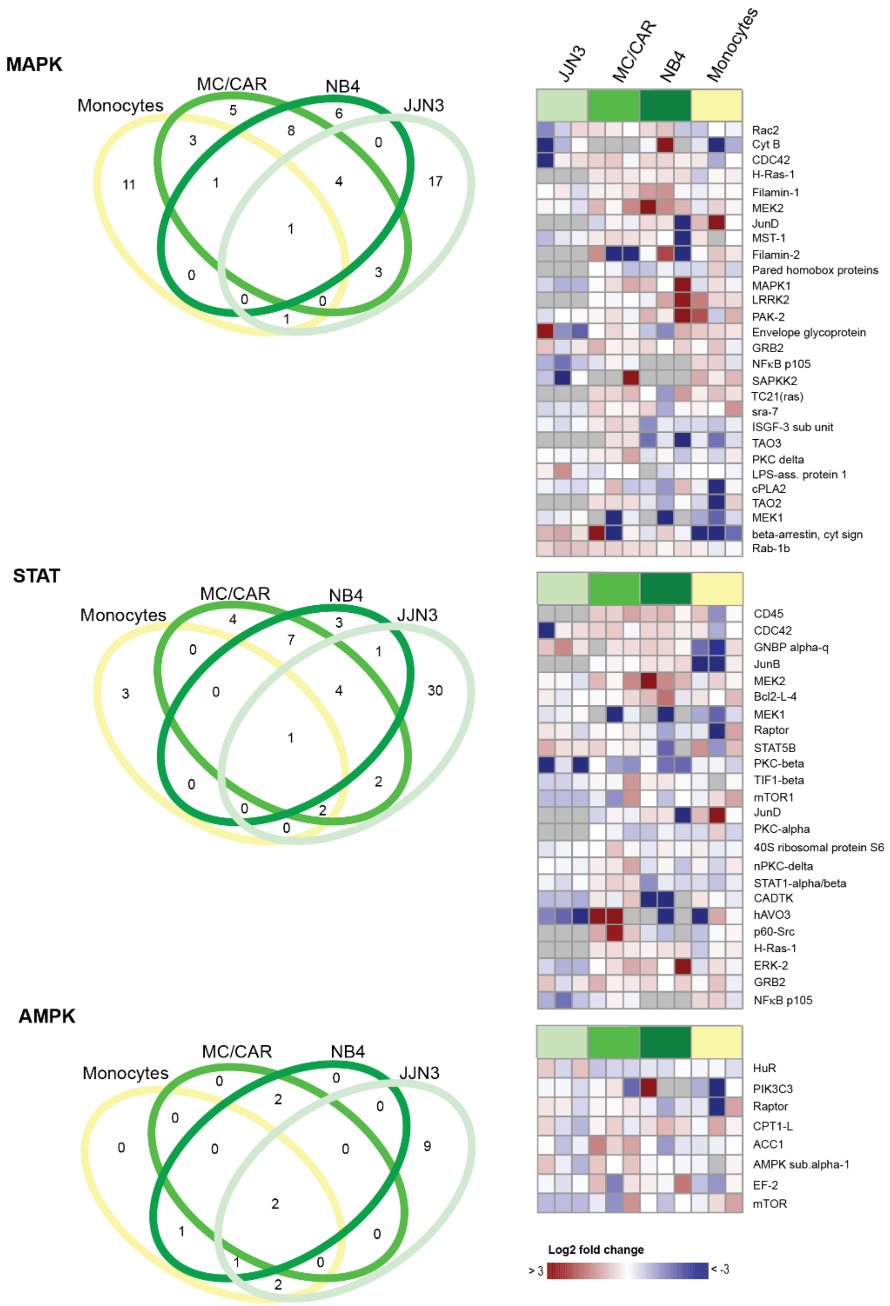


**Supplementary Figure S2. The signalome is strongly affected by targeting PCNA.** Results of MIB-assay of extract from haematological cells treated with APIM-peptide (JJN3: 6µM, MC/CAR, NB4, and primary monocytes: 8 µM) for 4h. Data from three repeated experiments is shown, each presented as $\log_2$ fold change relative to untreated control. Left panels: Venn diagrams displaying number of significantly changed proteins in MAPK, STAT and AMPK signalling pathways in APIM-peptide treated cells relative to untreated control in JJN3, MC/CAR and NB4 cells, and primary monocytes from three donors, according to the Wilcoxon Sign Rank test. Right panels: Heatmaps displaying proteins in the selected pathways that are significant changed from untreated control in at least two cell lines, according to the Wilcoxon Sign Rank test.

### Supplementary Figure S3, related to Figure 5

A

| Separation tecnique | Ionization | Metabolite coverage | Total detected ions | log2 > 1 / log2 < -1 | | |
|---|---|---|---|---|---|---|
| | | | | Cisplatin | APIM-peptide | APIM-peptide + cisplatin |
| capIC | ESI- | Anionic | 529 | 46 / 0 | 27 / 26 | 45 / 24 |
| HILIC | ESI- | Polar | 749 | 79 / 0 | 37 / 8 | 54 / 8 |
| RP | ESI- | Non to medium polar | 506 | 12 / 0 | 10 / 1 | 23 / 5 |
| RP | ESI+ | Non to medium polar | 848 | 138 / 0 | 27 / 9 | 165 / 15 |

B

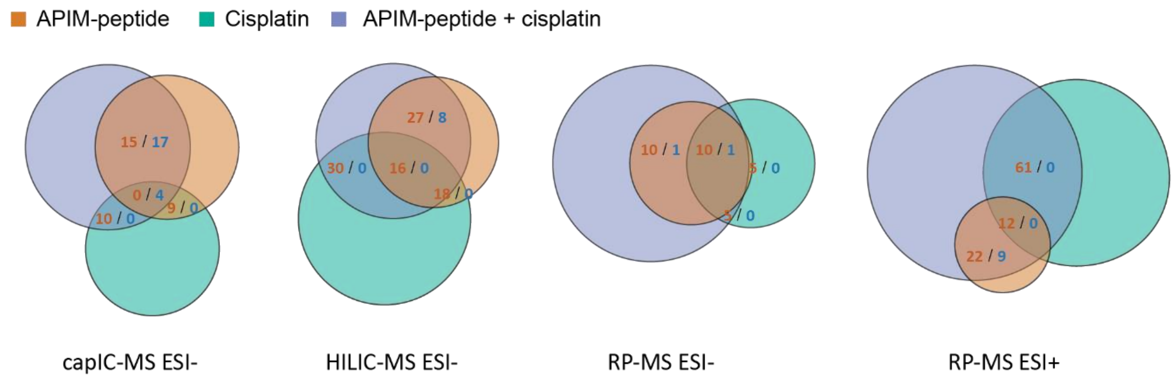


**Supplementary Figure S3. Targeting PCNA with the APIM-peptide alters the global metabolome of JJN3 cells.** Total number of up- (red) and downregulated (blue) metabolite pools in JJN3 cells 4h after treatment with APIM-peptide (8 µM), cisplatin (1 µM), or the combination as measured by capIC-MS ESI-, HILIC-MS ESI- , RP-MS ESI- and RP-MS ESI+ (**A**) out of total number of detected ions and (**B**) graphed as venn diagrams showing the number of up or down regulated ions common to the treatments, n=4. The threshold for up- and down-regulation is set to $\log_2$ fold change (treatment/control) > 1 and <-1, respectively. ESI: Electrospray ionization.

### Supplementary Figure S4, related to Figure 5

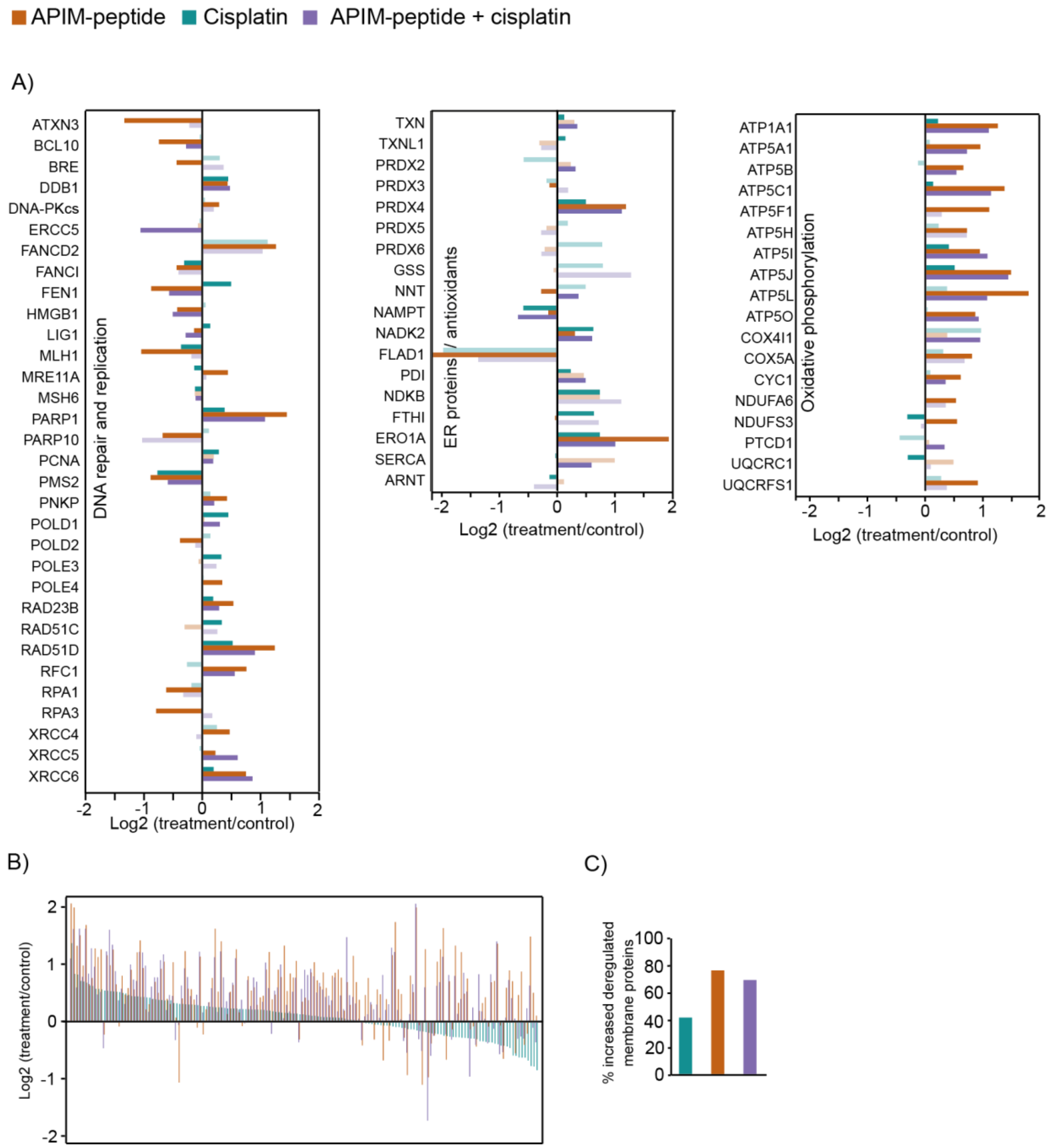


**Supplementary Figure S4: Treatment with APIM-peptide affects proteins regulating multiple other signaling pathways** (**A**) Results of MIB-assay of extract from JJN3 cells treated with APIM-peptide (6 µM), cisplatin (1 µM), or combination for 4h. Left panel: proteins involved in DNA repair and replication, mid panel: ER proteins and/or antioxidants, right panel: protein involved in oxidative phosphorylation. Data shown is mean from three repeated experiments, presented as $\log_2$ fold change relative to untreated control. Dark colour bars indicate significant change from untreated control according to the Wilcoxon Sign Rank test. (**B**) Transmembrane proteins detected by the MIB-assay. Proteins significantly changed from untreated control in at least one treatment according to the Wilcoxon Sign Rank test are shown. P-value, paired t-test: APIM-peptide vs cisplatin: $4.57*10^{-18}$. (**C**) Percent of upregulated transmembrane proteins relative to all deregulated transmembrane proteins detected by the MIB-assay.

Supplementary Figure S5, related to Figure 5

**A**

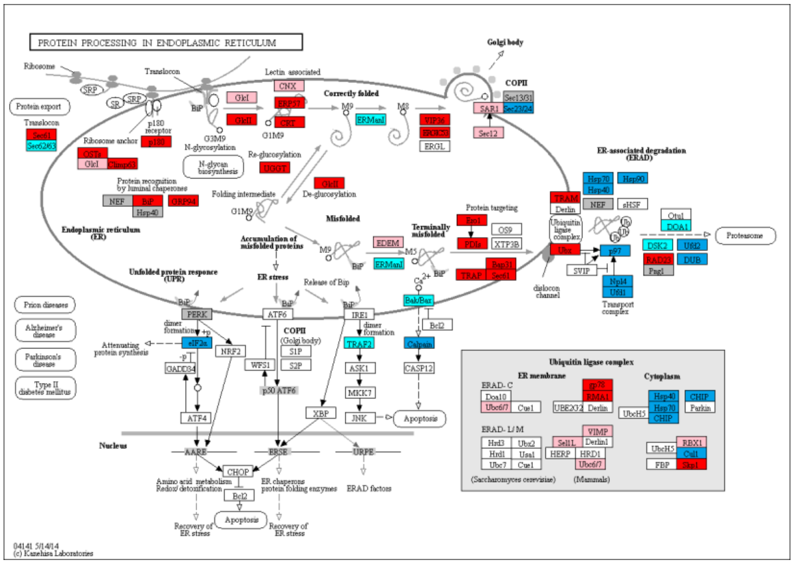

**B**

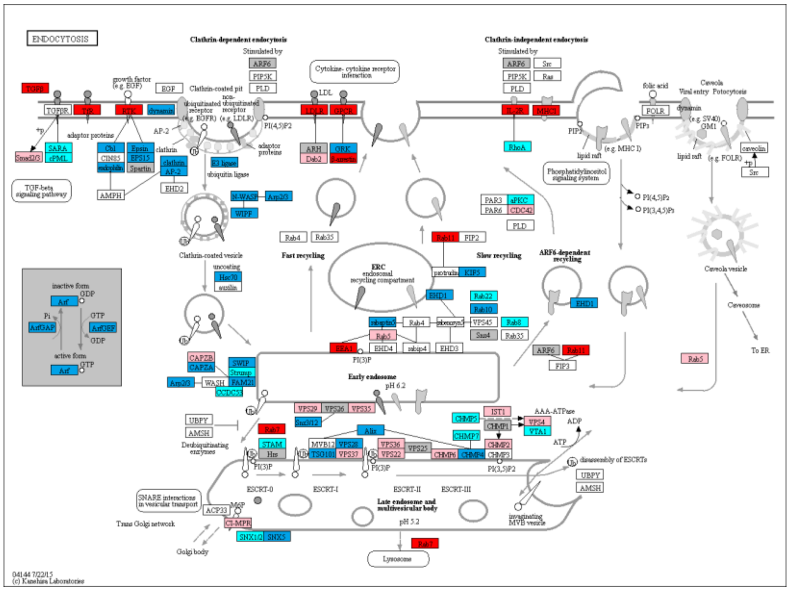

**Supplementary Figure S5. Deregulated pathways in JJN3 after APIM-peptide treatment**

Deregulated proteins detected by the MIB assay were subjected to KEGG pathway analysis for (**A**) Protein processing in ER and (**B**) Endocytosis. Red boxes indicate increased, blue boxes indicate decreased, and light colour indicates no significant change in proteins compared to control according to the Wilcoxon Sign Rank test. Grey boxes mean no change from control and white boxes means no data.

### Supplementary Figure S6, related to Figure 5

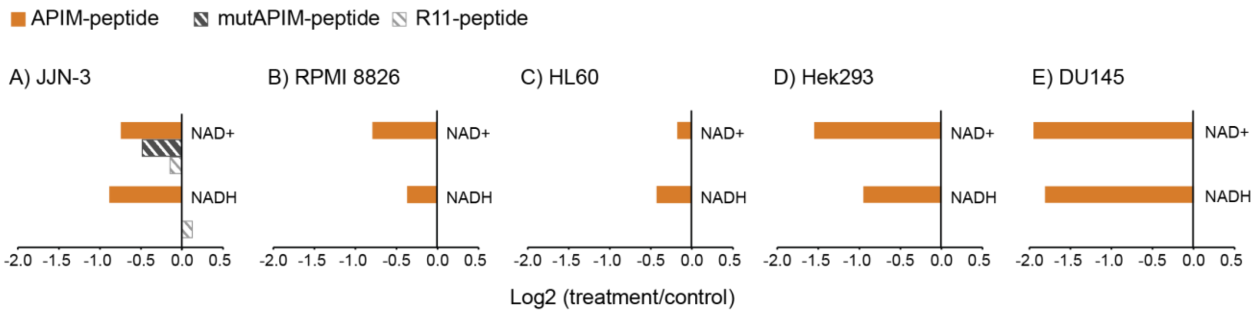


**Supplementary Figure S6: Treatment with APIM-peptide alter intracellular NAD levels.** $Log_2$ fold change of average intracellular $NAD^+$ and NADH levels in (**A**) APIM-peptide, mutAPIM-peptide and R11-peptide treated (8 µM) JJN3 cells , and APIM-peptide treated (8 µM) (**B**) RPMI 8226, (**C**) HL60, (**D**) Hek293, (**E**) DU145 cells relative to untreated control after 4h, n ≥ 4.

**Supplementary Table S1A:** Metabolite abbreviations and HMDB ID's

| Abbreviation | Common name | HMDB ID |
|---|---|---|
| 2-/3PG | 2-Phospho-D-glycerate | HMDB0000807 |
| | 3-Phospho-D-glycerate | HMDB0000362 |
| 2HG | 2-Hydroxyglutarate | HMDB0059655 |
| 6PG | 6-Phospho-D-gluconate | HMDB0001316 |
| ADP | Adenosine diphosphate | HMDB0001341 |
| aKG | a-Ketoglutaric acid | HMDB0000208 |
| Ala | L-Alanine | HMDB0000161 |
| AMP | Adenosine monophosphate | HMDB0000045 |
| Arg | L-Arginine | HMDB0000517 |
| Asn | L-Asparagine | HMDB0000168 |
| Asp | L-Aspartic acid | HMDB0000191 |
| ATP | Adenosine triphosphate | HMDB0000538 |
| cAMP | Cyclic AMP | HMDB0000058 |
| CDP | Cytidine diphosphate | HMDB0001546 |
| cGMP | Cyclic GMP | HMDB0001314 |
| Cit | Citric acid | HMDB0000094 |
| CMP | Cytidine monophosphate | HMDB0000095 |
| CTP | Cytidine triphosphate | HMDB0000082 |
| Cys | L-Cysteine | HMDB0000574 |
| dADP | Deoxyadenosine diphosphate | HMDB0001508 |
| dAMP | Deoxyadenosine monophosphate | HMDB0000905 |
| dATP | Deoxyadenosine triphosphate | HMDB0001532 |
| dCTP | Deoxycytidine triphosphate | HMDB0000998 |
| dGDP | Deoxyguanosine diphosphate | HMDB0000960 |
| dGMP | Deoxyguanosine monophosphate | HMDB0001044 |
| dGTP | Deoxyguanosine triphosphate | HMDB0001440 |
| dTDP | Thiamine diphosphate | HMDB0001274 |
| dTMP | Thiamine monophosphate | HMDB0002666 |
| dTTP | Thymidine triphosphate | HMDB0001342 |
| dUMP | Deoxyuridine monophosphate | HMDB0001409 |
| dUTP | Deoxyuridine triphosphate | HMDB0001191 |
| F1,6BP | Fructose 1,6-bisphosphate, | HMDB0001058 |
| F1P | Fructose 1-phosphate | HMDB0001076 |
| F6P | Fructose 6-phosphate | HMDB0000124 |
| Fum | Fumaric acid | HMDB0000134 |
| G1P/M1P | Glucose 1-phosphate | HMDB0001586 |
| | Mannose 1-phosphate | HMDB0006330 |
| G6P | Glucose 6-phosphate | HMDB0001401 |
| GA3P | Glyceraldehyde 3-phosphate | HMDB0001112 |
| GAL1P | Galactose 1-phosphate | HMDB0000645 |
| GAL6P | Galactose 6-phosphate | - |
| GDP | Guanosine diphosphate | HMDB0001201 |
| GL3P | Glycerol 3-phosphate | HMDB0000126 |
| Gln | L-Glutamine | HMDB0000641 |
| Glu | L-Glutamic acid | HMDB0000148 |
| Gly | Glycine | HMDB0000123 |
| GMP | Guanosine monophosphate | HMDB0001397 |
| GTP | Guanosine triphosphate | HMDB0001273 |
| His | L-Histidine | HMDB0000177 |
| ICit | Isocitric acid | HMDB0000193 |
| Ile | L-Isoleucine | HMDB0000172 |
| Lac | L-Lactic acid | HMDB0000190 |
| Leu | L-Leucine | HMDB0000687 |
| Lys | L-Lysine | HMDB0000182 |
| M6P | Mannose 6-phosphate | HMDB0001078 |
| Mal | L-Malic acid | HMDB0000156 |
| Met | L-Methionine | HMDB0000696 |
| $NAD^+$ | Nicotinamide Adenine Dinucleotide oxidized | HMDB0000902 |
| NADH | Nicotinamide Adenine Dinucleotide reduced | HMDB0001487 |
| Orn | L-ornithine | HMDB0000214 |
| PEP | Phosphoenolpyruvic acid | HMDB0000263 |
| Phe | L-Phenylalanine | HMDB0000159 |
| Pro | L-Proline | HMDB0000162 |
| PRPP | Phosphoribosyl pyrophosphate | HMDB0000280 |
| Pyr | Pyruvic acid | HMDB0000243 |
| R5P | D-Ribose 5-phosphate | HMDB0001548 |
| S7P | D-Sedoheptulose 7-phosphate | HMDB0001068 |
| Ser | L-Serine | HMDB0000187 |
| Suc | Succinic acid | HMDB0000254 |
| Thr | L-Threonine | HMDB0000167 |
| Trp | L-Tryptophan | HMDB0000929 |
| Tyr | L-Tyrosine | HMDB0000158 |
| UDP | Uridine diphosphate | HMDB0000295 |
| UDP-Glc/Gal-NAc | Uridine diphosphate-N-acetylglucosamine | HMDB0000290 |
| | Uridine diphosphate-N-acetylgalactosamine | HMDB0000304 |
| UDP-glu/gal | Uridine diphosphate glucose | HMDB0000286 |
| | Uridine diphosphategalactose | HMDB0000302 |
| UMP | Uridine monophosphate | HMDB0000288 |
| UTP | Uridine triphosphate | HMDB0000285 |
| Val | L-Valine | HMDB0000883 |

**Supplementary Table S1B:** $Log_2$ fold change of all measured central carbon metabolites in APIM-peptide treated (8 µM) cells relative to untreated control cells listed for JJN-3, RPMI 8226, MC/CAR, HL60, NB4, primary monocytes, DU145 and Hek293. Each of n ≥ 3 from one (RPMI8226, HL60, T24, DU145, and Hek293), or the average of n ≥ 3 from three (JJN3, MC/CAR, NB4, monocytes) repeated experiments is shown. Details on replicate numbers are listed in Supplementary Table S2. Metabolite abbreviations are listed in Supplementary Table S1A.

| Class | Metabolite | JJN3 | | | RPMI 8226 | | | MC/CAR | | | HL60 | | | | | NB-4 | | | Primary monocytes | | | DU145 | | | Hek293 | | |
|---|---|---|---|---|---|---|---|---|---|---|---|---|---|---|---|---|---|---|---|---|---|---|---|---|---|---|---|
| Glycolysis, PPP and other phosphorylated sugars | G6P | -1.5 | -1.0 | -1.6 | -0.5 | -1.3 | -1.1 | -0.5 | -0.2 | -0.1 | -1.3 | -1.8 | -0.8 | -1.0 | -1.5 | -0.7 | -0.9 | -1.2 | 0.0 | 0.1 | 0.0 | -0.2 | -0.3 | -0.3 | -0.2 | -0.1 | 0.2 |
| | F6P | -1.5 | -2.1 | -1.6 | -0.1 | -1.2 | -0.9 | -0.5 | -0.2 | 0.0 | -0.9 | -1.8 | -0.7 | -1.2 | -1.6 | -0.7 | -0.8 | -1.2 | -0.1 | 0.1 | 0.0 | 0.0 | -0.1 | -0.1 | -0.3 | 0.0 | 0.2 |
| | M6P | -1.5 | -1.1 | -1.3 | -0.2 | -1.2 | -1.1 | -0.5 | -0.2 | -0.2 | -1.0 | -1.6 | -0.7 | -1.1 | -1.6 | -0.6 | -0.9 | -1.2 | -0.1 | 0.1 | 0.1 | -0.6 | -0.3 | -0.3 | -0.3 | 0.1 | -0.4 |
| | GAL1P | -0.9 | NA | NA | -0.2 | -1.0 | -0.7 | -1.0 | 0.1 | -0.5 | -0.1 | -1.2 | -0.4 | -0.8 | -1.0 | -0.4 | 0.2 | -1.1 | -0.4 | -0.1 | 0.4 | NA | NA | NA | NA | NA | NA |
| | G-/M1P | -1.1 | NA | NA | -0.4 | -1.3 | -1.0 | -0.4 | -0.1 | -0.4 | -1.4 | -1.9 | -0.6 | -1.0 | -1.4 | -0.6 | -0.9 | -1.1 | -0.2 | 0.1 | 0.6 | 0.0 | -0.3 | -0.2 | -0.1 | 0.2 | -0.2 |
| | F1P | NA | NA | NA | -0.6 | -1.7 | -1.6 | -0.6 | -0.3 | 0.2 | -0.7 | -1.6 | 0.1 | -0.4 | -1.5 | 1.0 | 0.1 | -0.3 | 0.3 | 0.9 | 1.4 | 0.3 | 0.0 | 0.5 | -0.3 | 0.3 | -0.3 |
| | F1,6BP | NA | -1.8 | -0.8 | NA | NA | NA | 0.6 | -0.6 | -0.1 | NA | NA | NA | NA | NA | -0.5 | -0.9 | -0.8 | 0.1 | 0.7 | 0.6 | -0.4 | 0.2 | 0.1 | 0.5 | -0.4 | 0.7 |
| | GL3P | -0.7 | -1.9 | -1.6 | -0.9 | -1.6 | -1.4 | -0.4 | 0.0 | -0.2 | -2.0 | -2.0 | -0.7 | -0.6 | -1.1 | -0.4 | -0.6 | -0.9 | -0.2 | 0.1 | -0.2 | 0.1 | -0.9 | 0.6 | -0.6 | 0.1 | 0.0 |
| | 2/-3PG | -0.8 | -0.9 | -1.1 | -1.4 | -2.3 | -1.4 | -0.5 | -0.3 | -0.4 | -2.7 | -2.6 | -0.6 | -1.4 | -2.4 | -0.6 | -0.9 | -1.1 | 0.1 | 0.4 | 0.5 | -0.1 | 0.1 | 0.6 | -0.3 | 0.3 | 0.4 |
| | PEP | NA | -0.6 | -1.2 | NA | NA | NA | -0.6 | -0.4 | -0.4 | NA | NA | NA | NA | NA | -0.7 | -1.1 | -1.1 | -0.3 | 0.0 | 0.6 | NA | NA | 1.1 | -0.4 | 0.3 | 0.4 |
| | 6PG | -1.2 | -1.3 | -1.2 | -0.5 | -1.6 | -1.2 | -0.5 | -0.2 | -0.4 | -0.9 | -1.3 | -0.8 | -0.8 | -1.4 | -0.7 | -1.0 | -1.2 | 0.3 | 0.8 | 1.3 | 0.1 | 0.4 | 0.0 | 0.1 | -0.1 | 0.3 |
| | R5P | NA | NA | NA | NA | NA | NA | -0.4 | 0.0 | -0.5 | NA | NA | NA | NA | NA | -0.7 | -1.0 | -1.3 | 0.5 | 0.9 | 1.4 | -0.3 | -0.2 | -0.3 | -0.1 | 0.4 | 0.1 |
| | S7P | -2.4 | -1.8 | NA | -2.4 | NA | -2.3 | -0.4 | 0.0 | -0.3 | -2.2 | -2.9 | -0.8 | -1.3 | -2.0 | -0.5 | -0.7 | -1.0 | -0.2 | 0.0 | 0.7 | NA | NA | NA | NA | NA | NA |
| | PRPP | -1.2 | -2.1 | NA | -1.3 | -1.9 | -1.6 | -0.3 | -0.3 | -0.3 | -0.4 | -0.6 | -0.4 | -0.5 | -0.8 | -0.4 | -1.4 | -1.4 | -0.4 | 0.1 | 0.3 | NA | NA | 1.5 | 0.4 | 0.7 | -1.1 |
| (deoxy) Nucleoside phosphates | AMP | -0.2 | 0.7 | -0.4 | -0.4 | -0.5 | -0.8 | -0.5 | -0.1 | -0.5 | -1.9 | NA | -0.8 | -1.1 | -1.6 | -0.4 | -0.5 | -0.8 | -0.2 | -0.1 | -0.4 | 0.6 | -0.7 | 0.3 | -0.4 | -0.2 | -0.7 |
| | ADP | -0.7 | -0.4 | -1.1 | -0.6 | -1.2 | -0.9 | -0.6 | -0.1 | -0.5 | -2.0 | -2.6 | -1.0 | -1.4 | -1.9 | -0.3 | -0.5 | -0.9 | -0.4 | -0.1 | -0.4 | 0.8 | -1.3 | 0.0 | 0.3 | -0.1 | -0.3 |
| | ATP | -0.9 | -1.4 | -1.6 | -1.1 | -1.6 | -1.0 | -0.6 | -0.2 | -0.5 | -2.3 | -2.7 | -0.9 | -1.4 | -1.9 | -0.3 | -0.5 | -0.9 | -0.6 | -0.3 | -0.6 | -0.1 | 0.1 | -0.1 | 0.1 | 0.0 | 0.0 |
| | GMP | -0.3 | 0.0 | -0.6 | -0.8 | 0.0 | -0.6 | -0.3 | 0.0 | -0.4 | NA | 0.8 | -0.6 | -0.6 | -0.7 | -0.1 | -0.2 | -0.2 | -0.2 | -0.3 | 2.3 | -0.2 | -0.5 | 0.1 | 0.0 | 0.0 | -0.1 |
| | GDP | -0.9 | -0.2 | -1.2 | -0.3 | -1.4 | -1.4 | -0.6 | -0.2 | -0.5 | -1.6 | -3.8 | -1.4 | -2.5 | -3.8 | -0.2 | -0.4 | -0.8 | -0.3 | -0.1 | -0.3 | 0.6 | -0.6 | 0.3 | -0.2 | 0.2 | -0.5 |
| | GTP | -0.9 | -1.2 | -1.7 | -1.1 | -1.7 | -1.2 | -0.6 | -0.2 | -0.5 | -2.5 | -3.3 | -1.1 | -1.6 | -2.5 | -0.3 | -0.5 | -0.8 | -0.4 | -0.2 | -0.5 | 0.0 | 0.0 | -0.1 | -0.1 | -0.1 | 0.2 |
| | CMP | NA | -0.1 | NA | -1.2 | NA | NA | -0.3 | -0.1 | -0.3 | -3.2 | NA | 0.9 | NA | NA | 0.0 | -0.4 | -0.7 | 0.1 | 0.1 | 0.2 | NA | NA | NA | NA | NA | NA |
| | CDP | -1.0 | -0.2 | NA | -0.5 | -1.4 | -1.0 | -0.4 | -0.1 | -0.4 | -2.1 | -2.9 | -1.4 | -1.5 | -2.6 | -0.1 | -0.4 | -0.7 | -0.1 | 0.0 | -0.2 | 0.8 | -0.6 | 0.3 | 0.5 | 0.7 | -0.6 |
| | CTP | -1.3 | -1.4 | -1.5 | -1.3 | -2.0 | -1.3 | -0.6 | -0.2 | -0.4 | -2.7 | -3.4 | -1.1 | -1.6 | -2.3 | -0.3 | -0.5 | -0.9 | -0.3 | -0.1 | -0.5 | 0.0 | 0.1 | 0.1 | -0.1 | 0.2 | 0.1 |
| | UMP | -0.6 | -0.5 | -1.1 | -0.7 | -1.5 | -1.3 | -0.5 | -0.1 | -0.3 | -1.6 | -2.1 | -0.5 | -0.8 | -1.6 | -0.2 | -0.5 | -0.7 | -0.3 | -0.1 | -0.3 | 0.1 | -0.3 | 0.4 | -0.2 | 0.1 | 0.2 |
| | UDP | -1.4 | -0.3 | -1.2 | -0.9 | -1.5 | -1.1 | -0.4 | -0.1 | -0.4 | -2.4 | -4.9 | -1.3 | -2.3 | -2.9 | -0.3 | -0.4 | -0.9 | -0.4 | -0.1 | -0.3 | 1.2 | -1.2 | 0.4 | 0.7 | 0.6 | -0.8 |
| | UTP | -1.4 | -1.4 | -1.6 | -1.6 | -2.1 | -1.3 | -0.5 | -0.1 | -0.4 | -2.7 | -3.2 | -1.0 | -1.5 | -2.2 | -0.4 | -0.5 | -0.9 | -0.6 | -0.2 | -0.6 | NA | NA | 0.2 | 0.5 | -0.1 | -0.4 |
| | UDP-glu/-gal | NA | NA | NA | NA | NA | NA | -0.5 | -0.2 | -0.4 | NA | NA | NA | NA | NA | -0.3 | -0.5 | -0.7 | -0.7 | -0.1 | -0.4 | NA | NA | NA | NA | NA | NA |
| | UDP-Glc-NAc | -0.5 | -0.8 | -1.5 | -1.2 | -1.9 | -1.3 | -0.6 | -0.1 | -0.3 | -2.0 | -2.5 | -0.4 | -0.8 | -1.7 | -0.3 | -0.6 | -0.7 | -0.3 | 0.0 | -0.2 | 0.0 | 0.0 | 0.0 | -0.2 | 0.1 | -0.2 |
| | dAMP | NA | 0.1 | -0.6 | NA | NA | NA | NA | NA | NA | NA | NA | NA | NA | NA | NA | NA | NA | NA | NA | NA | 0.2 | NA | -2.1 | NA | NA | NA |
| | dADP | -1.1 | 0.1 | NA | 0.6 | -0.6 | -0.7 | -0.6 | -0.3 | -0.7 | 0.6 | -0.7 | -1.3 | -2.0 | -2.5 | -0.2 | -0.4 | -0.7 | NA | NA | NA | 0.9 | -1.1 | 0.1 | 0.4 | 0.4 | -1.0 |
| | dATP | -1.2 | -1.2 | -1.2 | -0.6 | -2.0 | -1.3 | -0.6 | -0.2 | -0.4 | -1.1 | -2.6 | -1.2 | -3.4 | -3.5 | -0.3 | -0.4 | -0.8 | NA | NA | NA | -0.2 | 0.2 | -0.2 | 0.1 | -0.2 | -0.2 |
| | dGDP | NA | -1.5 | -1.0 | NA | NA | NA | NA | NA | NA | NA | NA | NA | NA | NA | NA | NA | NA | NA | NA | NA | NA | NA | NA | NA | NA | NA |
| | dGTP | -1.1 | -0.9 | NA | 0.2 | -1.3 | -1.6 | -0.7 | -0.2 | -0.6 | 0.7 | -1.4 | -1.9 | NA | NA | -0.5 | -0.5 | -0.8 | NA | NA | NA | NA | NA | NA | NA | 1.5 | -0.4 |
| | dCTP | -1.0 | -0.7 | -0.6 | -0.6 | -1.1 | -0.9 | -0.5 | -0.1 | -0.4 | 0.1 | -2.5 | -1.5 | -2.5 | -2.7 | -0.3 | -0.5 | -0.8 | NA | NA | NA | -0.2 | 0.1 | 0.2 | 0.0 | -0.2 | 0.1 |
| | dUMP | -0.4 | NA | NA | 0.5 | -1.4 | -1.7 | NA | NA | NA | 1.1 | -1.1 | NA | -2.1 | NA | NA | NA | NA | NA | NA | NA | 0.0 | -1.1 | -0.3 | -0.4 | -0.4 | -0.1 |
| | dTMP | -0.2 | -0.2 | NA | 0.6 | -0.3 | -0.4 | -0.2 | -0.1 | -0.1 | 0.5 | -0.2 | -0.4 | -0.5 | -0.6 | -0.5 | -0.5 | -0.5 | NA | NA | NA | 0.1 | -0.1 | -0.1 | -0.4 | 0.0 | 0.0 |
| | dTDP | -0.7 | -0.1 | -0.5 | 0.2 | -0.6 | -0.6 | NA | NA | NA | 0.3 | -0.5 | -0.6 | -0.6 | -0.8 | -0.3 | -0.4 | -0.9 | NA | NA | NA | 1.0 | -1.1 | 0.4 | 0.6 | 0.7 | -0.6 |
| | dTTP | -1.6 | -1.2 | -1.0 | -1.1 | -3.0 | -1.6 | -0.7 | -0.1 | -0.7 | -0.8 | -6.1 | -2.0 | -3.5 | -5.3 | -0.3 | -0.5 | -1.0 | NA | NA | NA | 0.0 | 0.2 | 0.2 | -0.1 | 0.0 | -0.9 |
| TCA cycle and asisociated | Lac | NA | NA | NA | NA | NA | NA | -0.6 | -0.3 | -0.1 | NA | NA | NA | NA | NA | -0.4 | -0.3 | -0.5 | -0.8 | 0.2 | -0.5 | NA | 0.1 | -0.1 | 0.0 | 0.0 | -0.2 |
| | Pyr | NA | NA | NA | NA | NA | NA | -0.4 | -0.4 | -0.2 | NA | NA | NA | NA | NA | -0.4 | -0.4 | -0.5 | -1.9 | -0.1 | -1.0 | NA | -0.1 | -0.6 | 0.1 | 0.0 | 0.2 |
| | Cit | -0.7 | -1.4 | -0.9 | -0.4 | -1.2 | -1.3 | -0.5 | -0.3 | -0.2 | -0.5 | -0.9 | -0.2 | -0.3 | -0.7 | -0.1 | -0.2 | -0.3 | -0.3 | 0.0 | -0.8 | 0.0 | 0.0 | 0.2 | -0.3 | 0.1 | 0.0 |
| | Icit | -1.0 | -0.8 | -0.8 | -0.4 | -1.5 | -0.6 | NA | NA | NA | 0.2 | -0.5 | 0.3 | -0.2 | -1.1 | NA | NA | NA | NA | NA | NA | 0.1 | 0.0 | 0.2 | -0.2 | -0.1 | 0.0 |
| | aKG | 0.5 | -0.7 | -1.2 | 0.6 | 0.1 | 0.2 | -0.6 | -0.4 | -0.6 | 1.4 | 0.6 | 0.4 | 1.0 | 1.3 | -0.9 | -1.2 | -1.6 | -3.4 | 0.9 | -1.0 | NA | NA | NA | NA | NA | NA |
| | 2HG | 0.0 | 0.4 | -0.6 | 0.1 | -0.3 | -0.2 | NA | NA | NA | 0.0 | 0.0 | 0.2 | 0.2 | -0.2 | NA | NA | NA | -0.1 | 0.0 | -0.1 | -0.9 | -0.4 | -0.1 | -0.3 | 0.2 | 0.0 |
| | Suc | -0.2 | NA | -0.4 | 0.2 | -0.5 | -1.0 | -0.6 | -0.2 | -0.3 | -0.4 | -0.4 | -0.2 | 0.0 | -0.6 | -0.2 | -0.3 | -0.6 | 1.0 | 0.1 | -0.5 | -0.8 | 0.0 | 0.3 | -0.7 | 0.3 | 0.4 |
| | Fum | NA | NA | -0.5 | NA | NA | NA | -1.1 | -0.2 | -1.4 | NA | NA | NA | NA | NA | -0.8 | -0.7 | -0.7 | 0.4 | 0.2 | 0.6 | NA | -0.3 | -0.2 | -0.1 | 0.0 | 0.2 |
| | Mal | -0.7 | -0.7 | -1.2 | -0.5 | -1.1 | -0.8 | -0.5 | -0.3 | -0.4 | -0.3 | -0.3 | 0.2 | 0.3 | -0.3 | -0.3 | -0.5 | -0.8 | -0.4 | -0.3 | -0.1 | 0.1 | -0.2 | 0.2 | -0.2 | 0.1 | 0.0 |
| Amino acids | Ala | -0.6 | -0.5 | -1.6 | -0.1 | -0.3 | -0.2 | -0.7 | -0.6 | -0.3 | 0.0 | -0.8 | -0.4 | -0.2 | -0.3 | -0.4 | -0.3 | -0.7 | -0.2 | 0.1 | -0.1 | NA | 0.4 | 0.5 | 0.0 | 0.1 | 0.0 |
| | Arg | 0.4 | -1.1 | 0.2 | 0.4 | 0.5 | 0.4 | -0.5 | -0.5 | -0.1 | 0.1 | -0.5 | -0.4 | 0.1 | 0.1 | -0.4 | -0.2 | -0.6 | 0.2 | 0.1 | -0.1 | NA | NA | NA | NA | NA | NA |
| | Asn | -0.1 | -0.6 | -0.5 | -0.2 | -0.3 | -0.3 | -0.6 | -0.5 | -0.3 | -0.4 | -1.0 | -0.6 | -0.4 | -0.3 | -0.4 | -0.3 | -0.7 | 0.0 | 0.1 | -0.1 | NA | NA | NA | NA | NA | NA |
| | Asp | -1.2 | -1.6 | -2.7 | -0.3 | -0.3 | -0.2 | -0.6 | -0.4 | -0.4 | -0.1 | -0.8 | -0.6 | -0.2 | -0.2 | -0.2 | -0.1 | -0.4 | -0.3 | 0.1 | -0.1 | NA | 0.0 | 0.2 | 0.0 | 0.1 | 0.3 |
| | Cys | NA | 1.6 | 1.8 | NA | NA | NA | NA | NA | NA | NA | NA | NA | NA | NA | NA | NA | NA | NA | NA | NA | NA | NA | NA | NA | NA | NA |
| | Gln | -0.1 | -0.4 | -3.7 | -0.2 | 0.0 | -0.1 | -0.6 | -0.6 | -0.3 | -0.4 | -0.9 | -0.7 | -0.3 | -0.3 | -0.4 | -0.2 | -0.7 | 0.1 | 0.1 | -0.2 | NA | NA | NA | NA | NA | NA |
| | Glu | -1.1 | -1.5 | -3.1 | -0.7 | -0.7 | -0.7 | -0.7 | -0.4 | -0.4 | 0.7 | -1.2 | -0.6 | -1.3 | -0.6 | -0.4 | -0.4 | -0.8 | -0.2 | 0.0 | -0.2 | NA | -0.1 | 0.0 | 0.0 | 0.0 | 0.3 |
| | Gly | -1.0 | -0.3 | -1.5 | -0.4 | -0.6 | -0.5 | -0.6 | -0.5 | -0.3 | -0.7 | -1.3 | -1.0 | -0.9 | -0.8 | -0.4 | -0.4 | -0.5 | -0.3 | 0.0 | -0.2 | NA | NA | NA | NA | NA | NA |
| | His | -0.5 | -0.6 | -1.0 | 0.2 | 0.0 | -0.1 | -0.6 | -0.6 | -0.3 | 0.1 | -0.6 | -0.4 | 0.0 | -0.1 | -0.4 | -0.2 | -0.4 | 0.0 | 0.1 | -0.1 | NA | NA | NA | NA | NA | NA |
| | Ile | 0.0 | 0.3 | -0.8 | 0.2 | 0.1 | 0.1 | -0.6 | -0.6 | -0.2 | 0.0 | -0.6 | -0.5 | 0.1 | -0.1 | -0.4 | -0.2 | -0.6 | 0.0 | 0.1 | -0.1 | NA | NA | NA | NA | NA | NA |
| | Leu | 0.0 | 0.3 | -0.9 | 0.2 | 0.1 | 0.0 | -0.6 | -0.6 | -0.2 | 0.1 | -0.6 | -0.5 | 0.0 | -0.1 | -0.4 | -0.2 | -0.6 | 0.0 | 0.1 | -0.1 | NA | NA | NA | NA | NA | NA |
| | Lys | 0.1 | -0.4 | -0.7 | 0.4 | 0.5 | 0.3 | -0.6 | -0.6 | -0.2 | 0.3 | -0.4 | -0.4 | 0.2 | 0.1 | -0.3 | 0.0 | -0.4 | 0.1 | 0.1 | -0.1 | NA | NA | NA | NA | NA | NA |
| | Met | 0.7 | -0.8 | NA | -0.2 | 0.2 | 0.1 | -0.5 | -0.7 | -0.2 | -0.3 | -3.4 | -2.2 | -0.7 | -0.7 | -0.4 | -0.2 | -0.7 | 0.0 | 0.0 | -0.2 | NA | NA | NA | NA | NA | NA |
| | Phe | -0.1 | -0.1 | -0.8 | 0.2 | 0.1 | 0.0 | -0.6 | -0.6 | -0.2 | 0.0 | -0.7 | -0.5 | 0.0 | -0.1 | -0.3 | -0.2 | -0.5 | 0.0 | 0.1 | -0.1 | NA | 0.5 | 0.7 | 0.0 | 0.1 | 0.2 |
| | Pro | -0.7 | -0.8 | -1.5 | -0.4 | -0.3 | -0.3 | -0.6 | -0.5 | -0.3 | -0.7 | -1.3 | -0.9 | -0.7 | -0.7 | -0.5 | -0.5 | -0.9 | 0.0 | 0.1 | -0.1 | NA | 0.3 | 0.2 | -0.2 | 0.0 | 0.0 |
| | Ser | -1.1 | -0.9 | -0.5 | 0.0 | -0.4 | -0.3 | -0.6 | -0.4 | -0.3 | 0.0 | -0.7 | -0.6 | -0.2 | -0.3 | -0.2 | -0.2 | 0.0 | -0.2 | 0.1 | -0.1 | NA | NA | NA | NA | NA | NA |
| | Thr | -0.5 | -0.6 | -0.6 | 0.0 | -0.2 | -0.2 | -0.6 | -0.5 | -0.3 | -0.2 | -0.9 | -0.5 | -0.3 | -0.3 | -0.4 | -0.2 | -0.5 | 0.0 | 0.1 | -0.2 | NA | 0.6 | 0.8 | 0.1 | 0.1 | 0.2 |
| | Trp | -0.1 | -0.1 | -0.5 | 0.4 | 0.3 | 0.2 | -0.6 | -0.7 | -0.2 | 0.1 | -0.6 | -0.4 | 0.0 | 0.1 | -0.2 | 0.0 | -0.4 | 0.2 | 0.0 | -0.3 | NA | NA | NA | NA | NA | NA |
| | Tyr | -0.2 | -0.3 | -0.7 | 0.1 | 0.1 | 0.1 | -0.6 | -0.6 | -0.2 | 0.0 | -0.6 | -0.4 | 0.0 | -0.1 | -0.4 | -0.2 | -0.5 | 0.1 | 0.1 | -0.2 | NA | 0.3 | 0.6 | -0.1 | 0.0 | 0.2 |
| | Val | -0.1 | 0.6 | -0.9 | 0.2 | 0.1 | 0.1 | -0.6 | -0.6 | -0.2 | 0.1 | -0.6 | -0.4 | 0.1 | 0.0 | -0.3 | -0.2 | -0.5 | 0.1 | 0.1 | -0.1 | NA | NA | NA | 0.1 | 0.1 | 0.2 |

**Supplementary Table S1C**: Intracellular metabolites measured in all cell types (left column). Significant diffrences (ANOVA, Tukets post hoc test, p = 0.05) between hematological cancer cell lines and the remaining panel are indicated. Metabolite abbreviations are listed in Supplementary Table S1A.

| **Metabolite** | **Hematological cancer cell line** | **Significant differences (p = 0.05)** | | | |
|---|---|---|---|---|---|
| | | Primary monocytes | T24 | DU145 | Hek293 |
| G6P | JJN-3 | x | x | x | x |
| | RPMI 8226 | x | | | x |
| | MC/CAR | | | | |
| | HL60 | x | x | x | x |
| | NB4 | x | | | x |
| F6P | JJN-3 | x | x | x | x |
| | RPMI 8226 | | | | |
| | MC/CAR | | | | |
| | HL60 | x | | x | x |
| | NB4 | x | | | |
| M6P | JJN-3 | x | | x | x |
| | RPMI 8226 | x | | | |
| | MC/CAR | | | | |
| | HL60 | x | x | x | x |
| | NB4 | x | | | |
| GL3P | JJN-3 | | | | |
| | RPMI 8226 | | | | |
| | MC/CAR | | | | |
| | HL60 | | | x | |
| | NB4 | | | | |
| 2-/3PG | JJN-3 | | | | |
| | RPMI 8226 | x | x | x | x |
| | MC/CAR | | | | |
| | HL60 | x | x | x | x |
| | NB4 | | | | |
| 2-/3PG | JJN-3 | | | | |
| | RPMI 8226 | x | x | x | x |
| | MC/CAR | | | | |
| | HL60 | x | x | x | x |
| | NB4 | | | | |
| 6PG | JJN-3 | x | x | x | x |
| | RPMI 8226 | x | x | x | x |
| | MC/CAR | x | | | |
| | HL60 | x | x | x | x |
| | NB4 | x | x | x | x |
| ADP | JJN-3 | | | | |
| | RPMI 8226 | | | | |
| | MC/CAR | | | | |
| | HL60 | x | x | x | x |
| | NB4 | | | | |
| ATP | JJN-3 | | x | x | x |
| | RPMI 8226 | | x | x | x |
| | MC/CAR | | | | |
| | HL60 | x | x | x | x |
| | NB4 | | | | |
| GTP | JJN-3 | | x | | x |
| | RPMI 8226 | | x | x | x |
| | MC/CAR | | | | |
| | HL60 | x | x | x | x |
| | NB4 | | | | |
| CTP | JJN-3 | | | x | x |
| | RPMI 8226 | | | x | x |
| | MC/CAR | | | | |
| | HL60 | x | x | x | x |
| | NB4 | | | | |
| GDP | JJN-3 | | | | |
| | RPMI 8226 | | | | |
| | MC/CAR | | | | |
| | HL60 | x | x | x | x |
| | NB4 | | | | |
| UMP | JJN-3 | | | | |
| | RPMI 8226 | | | x | x |
| | MC/CAR | | | | |
| | HL60 | x | x | x | x |
| | NB4 | | | | |
| UDP | JJN-3 | | | | |
| | RPMI 8226 | | | | |
| | MC/CAR | | | | |
| | HL60 | x | x | x | x |
| | NB4 | | | | |
| Cit | JJN-3 | | x | x | x |
| | RPMI 8226 | | x | x | x |
| | MC/CAR | | | | |
| | HL60 | | | | |
| | NB4 | | | | |
| Mal | JJN-3 | | x | x | x |
| | RPMI 8226 | | x | x | x |
| | MC/CAR | | | | |
| | HL60 | | | | |
| | NB4 | | | | |

**Supplementary Table S1D:** Heatmapped $log_2$ fold change of all measured central carbon metabolites in JJN3 cells treated with cisplatin (1 µM), APIM-peptide (8 µM), or the respective combination for 4, 8 and 24 hours relative to untreated control cells. Blue: descreased, red: increased. Metabolite abbreviations are listed in Supplementary Table S1A.

| Class | Metabolite | 4h | | | 8h | | | 24h | | |
|---|---|---|---|---|---|---|---|---|---|---|
| | | Cisplatin | APIM-peptide | APIM-peptide + cisplatin | Cisplatin | APIM-peptide | APIM-peptide + cisplatin | Cisplatin | APIM-peptide | APIM-peptide + cisplatin |
| Glycolysis, PPP and other phosphorylated sugars | G6P | -0.9 | -1.6 | -2.9 | -0.3 | 1.9 | 0.0 | -0.3 | -1.4 | -1.5 |
| | F6P | -0.9 | -1.6 | -3.2 | -0.9 | 2.5 | 0.2 | -0.5 | -1.6 | -1.8 |
| | M6P | -0.7 | -1.3 | -1.9 | -0.3 | 1.6 | -0.2 | -0.4 | -1.4 | -1.5 |
| | F1,6BP | -0.6 | -0.8 | -1.0 | -0.1 | 1.1 | 0.1 | -0.1 | -0.7 | -0.5 |
| | GL3P | -0.9 | -1.6 | -2.4 | -0.3 | 2.0 | 0.0 | -0.5 | -1.2 | -1.3 |
| | 2/-3PG | -0.8 | -1.1 | -2.3 | -0.2 | 1.7 | 0.0 | -0.6 | -1.7 | -1.9 |
| | PEP | -0.9 | -1.2 | -1.7 | 0.0 | 2.3 | 0.2 | -0.7 | -1.6 | -2.1 |
| (deoxy) Nucleoside phosphates | AMP | -0.6 | -0.4 | -0.8 | 0.3 | 0.7 | -0.9 | -0.1 | -0.5 | -0.7 |
| | ADP | -0.9 | -1.1 | -1.8 | -0.1 | 1.2 | 0.1 | -0.3 | -1.3 | -1.4 |
| | ATP | -0.9 | -1.6 | -2.7 | -0.6 | 1.6 | -0.4 | -0.4 | -2.1 | -2.2 |
| | GMP | -0.4 | -0.6 | -0.6 | 0.0 | 0.6 | 0.2 | 0.0 | -0.6 | -0.7 |
| | GDP | -0.9 | -1.2 | -1.7 | -0.1 | 1.1 | -0.1 | -0.4 | -1.4 | -1.7 |
| | GTP | -1.0 | -1.7 | -2.6 | -0.5 | 1.5 | -0.2 | -0.4 | -1.9 | -1.9 |
| | CTP | -0.9 | -1.5 | -2.4 | -0.4 | 1.7 | -0.1 | -0.4 | -1.0 | -0.8 |
| | UMP | -0.7 | -1.1 | -1.4 | -0.2 | 1.3 | 0.3 | -0.1 | -0.9 | -1.0 |
| | UDP | -1.0 | -1.2 | -1.7 | -0.1 | 1.5 | 0.1 | -0.4 | -1.2 | -1.6 |
| | UTP | -1.2 | -1.6 | -2.7 | -0.4 | 1.7 | -0.1 | -0.6 | -1.6 | -1.7 |
| | UDP-GlcNAc | -1.3 | -1.5 | -2.0 | -0.3 | 1.4 | -0.3 | -0.2 | -1.5 | -1.6 |
| | dADP | -0.6 | -0.6 | -0.8 | 0.2 | 1.0 | 0.3 | 1.3 | 0.5 | 0.6 |
| | dATP | -0.8 | -1.2 | -1.6 | -0.3 | 1.4 | -0.2 | -0.3 | -1.7 | -1.5 |
| | dGTP | -0.6 | -1.0 | -1.2 | -0.1 | 1.4 | 0.2 | -0.2 | -1.2 | -0.9 |
| | dCTP | -0.4 | -0.6 | -0.6 | 0.1 | 1.1 | 0.2 | 0.1 | -0.5 | -0.3 |
| | dTDP | -0.5 | -0.5 | -0.4 | 0.1 | 0.8 | 0.3 | 0.1 | -0.5 | -0.4 |
| | dTTP | -0.7 | -1.0 | -1.4 | 0.0 | 1.6 | 0.2 | -0.2 | -1.2 | -0.9 |
| TCA cycle and asisociated | Lac | 0.0 | -0.1 | 0.0 | 0.6 | 1.5 | 0.7 | 0.1 | -0.5 | -0.6 |
| | Pyr | -0.2 | -0.1 | 0.2 | 0.3 | 0.6 | 0.7 | 0.4 | 0.5 | 0.7 |
| | Cit | -0.6 | -0.9 | -0.9 | 0.1 | 1.0 | 0.1 | 0.0 | -1.2 | -1.2 |
| | Icit | -0.7 | -0.8 | -1.0 | 0.5 | 1.7 | 0.5 | 0.0 | -1.2 | -1.3 |
| | aKG | -0.7 | -1.2 | NA | -0.5 | 1.4 | -0.7 | -0.5 | -1.8 | -1.9 |
| | 2HG | -0.7 | -0.6 | -0.9 | -0.3 | 0.8 | 0.2 | -0.4 | -1.1 | -0.9 |
| | Suc | -0.6 | -0.4 | -0.2 | 0.1 | 0.3 | 0.4 | -0.1 | -0.4 | 0.1 |
| | Fum | -0.5 | -0.5 | -0.5 | -0.3 | 1.0 | 0.1 | -0.4 | -1.2 | -1.2 |
| | Mal | -0.9 | -1.2 | -1.8 | -0.4 | 1.5 | -0.1 | -0.4 | -1.6 | -1.6 |
| Amino acids | Ala | -0.3 | -1.6 | -0.8 | 0.3 | -0.4 | -0.1 | 1.4 | -1.0 | -0.9 |
| | Arg | 0.0 | 0.2 | -0.3 | 0.0 | 0.4 | -0.4 | 0.7 | 0.5 | 0.4 |
| | Asn | -0.2 | -0.5 | -0.7 | 0.1 | -0.6 | -0.5 | 0.6 | -1.1 | -1.2 |
| | Asp | -0.3 | -2.7 | -2.3 | -0.2 | -1.1 | -1.3 | -0.1 | -2.8 | -3.3 |
| | Cys | 0.8 | 1.8 | 4.0 | -2.3 | 1.7 | 2.3 | -0.8 | 2.6 | 4.0 |
| | Gln | 0.3 | -3.7 | 0.0 | 0.3 | -0.8 | -0.9 | -0.3 | -1.9 | 0.2 |
| | Glu | -0.2 | -3.1 | -2.1 | 0.2 | -1.0 | -1.0 | 0.6 | -2.2 | -2.6 |
| | Gly | -0.3 | -1.5 | -1.0 | 0.0 | -0.5 | -0.3 | 0.7 | -1.2 | -1.2 |
| | His | -0.3 | -1.0 | -0.7 | 0.0 | -0.3 | -0.5 | 0.2 | -1.0 | -0.8 |
| | Ile | -0.1 | -0.8 | 0.0 | 0.3 | 0.0 | 0.1 | 0.6 | -0.6 | -0.5 |
| | Leu | -0.2 | -0.9 | -0.3 | 0.1 | -0.2 | -0.2 | 0.2 | -1.0 | -0.8 |
| | Lys | -0.1 | -0.7 | -0.1 | 0.0 | 0.1 | 0.0 | -0.1 | -0.6 | -0.4 |
| | Phe | -0.2 | -0.8 | -0.3 | 0.1 | -0.2 | -0.2 | 0.2 | -1.0 | -0.9 |
| | Pro | -0.3 | -1.5 | -1.1 | 0.0 | -0.6 | -0.5 | 0.6 | -1.6 | -1.7 |
| | Ser | -0.2 | -0.5 | -0.6 | 0.2 | -0.1 | 0.0 | 0.6 | -0.5 | -0.7 |
| | Thr | -0.1 | -0.6 | -0.6 | 0.1 | -0.2 | -0.2 | 0.5 | -0.8 | -0.9 |
| | Trp | -0.4 | -0.5 | -0.1 | 0.2 | 0.3 | 0.0 | 0.3 | -0.3 | -0.1 |
| | Tyr | -0.1 | -0.7 | -0.3 | 0.2 | -0.1 | -0.1 | 0.3 | -0.9 | -0.7 |
| | Val | -0.2 | -0.9 | -0.2 | 0.1 | -0.1 | -0.1 | 0.2 | -0.9 | -0.8 |

**Supplementary Table S1E:** Heatmapped log2 fold change of all measured central carbon metabolites in DU145 cells treated with cisplatin (1 µM), APIM-peptide (8 µM), or the respective combination for 4, 8 and 24 hours relative to untreated control cells. Blue: descreased, red: increased. Metabolite abbreviations are listed in Supplementary Table S1A.

| Class | Metabolite | 4h | | | 8h | | | 24h | | |
|---|---|---|---|---|---|---|---|---|---|---|
| | | Cisplatin | APIM-peptide | APIM-peptide + cisplatin | Cisplatin | APIM-peptide | APIM-peptide + cisplatin | Cisplatin | APIM-peptide | APIM-peptide + cisplatin |
| Glycolysis, PPP and other phosphorylated sugars | G6P | 0.6 | -0.2 | -0.1 | -0.4 | -0.3 | -0.7 | 0.4 | -0.3 | 0.4 |
| | F6P | 0.6 | 0.0 | 0.0 | -0.4 | -0.1 | -0.8 | 0.4 | -0.1 | 0.3 |
| | M6P | 0.3 | -0.6 | -0.7 | -0.5 | -0.3 | -0.7 | 1.1 | -0.3 | 0.5 |
| | G-/M1P | 0.4 | 0.0 | 0.0 | -0.3 | -0.3 | -0.9 | 0.8 | -0.2 | 0.7 |
| | F1P | 0.6 | 0.3 | 0.8 | 0.2 | 0.0 | -0.2 | 0.2 | 0.5 | 0.4 |
| | F1,6BP | -0.5 | -0.4 | 0.1 | 0.8 | 0.2 | 0.0 | -1.5 | 0.1 | -0.5 |
| | GL3P | 0.0 | 0.1 | -0.3 | -0.1 | -0.9 | -0.8 | 0.8 | 0.6 | 1.2 |
| | 2-/3PG | 0.2 | -0.1 | -0.1 | 0.4 | 0.1 | 0.0 | 0.4 | 0.6 | 0.9 |
| | PEP | NA | NA | -1.0 | NA | NA | NA | NA | 1.1 | 2.8 |
| | 6PG | 0.0 | 0.1 | -0.2 | 0.2 | 0.4 | 0.4 | 0.3 | 0.0 | 0.0 |
| | R5P | 0.6 | -0.3 | 0.1 | 0.3 | -0.2 | -0.1 | 0.0 | -0.3 | 0.0 |
| | PRPP | 0.6 | NA | -0.3 | NA | NA | NA | 1.1 | 1.5 | 1.4 |
| (deoxy) Nucleoside phosphates | AMP | 0.1 | 0.6 | -0.1 | -0.7 | -0.7 | -0.2 | 0.7 | 0.3 | 0.9 |
| | ADP | 0.1 | 0.8 | 0.1 | -1.3 | -1.3 | -0.2 | 0.4 | 0.0 | 0.3 |
| | ATP | 0.0 | -0.1 | 0.0 | 0.1 | 0.1 | 0.0 | -0.1 | -0.1 | -0.1 |
| | GMP | -0.2 | -0.2 | 0.1 | 0.3 | -0.5 | -0.5 | 0.2 | 0.1 | 0.1 |
| | GDP | 0.1 | 0.6 | 0.2 | -0.5 | -0.6 | 0.0 | 0.4 | 0.3 | 0.3 |
| | GTP | 0.0 | 0.0 | 0.2 | 0.3 | 0.0 | 0.1 | 0.0 | -0.1 | 0.2 |
| | CDP | 0.2 | 0.8 | 0.3 | -0.5 | -0.6 | 0.2 | 0.3 | 0.3 | 0.1 |
| | CTP | -0.3 | 0.0 | -0.1 | 0.1 | 0.1 | -0.2 | -0.1 | 0.1 | 0.1 |
| | UMP | 0.0 | 0.1 | -0.2 | -0.3 | -0.3 | -0.1 | 0.1 | 0.4 | 0.3 |
| | UDP | 0.2 | 1.2 | 0.1 | -1.2 | -1.2 | 0.1 | 0.5 | 0.4 | 0.6 |
| | UTP | -2.6 | NA | -1.5 | NA | NA | NA | 0.7 | 0.2 | 1.2 |
| | UDP-Glc-NAc | 0.2 | 0.0 | 0.1 | 0.0 | 0.0 | 0.1 | 0.1 | 0.0 | -0.2 |
| | dAMP | 0.2 | 0.2 | 0.5 | NA | NA | NA | -3.9 | -2.1 | -0.5 |
| | dADP | 0.0 | 0.9 | 0.1 | -0.2 | -1.1 | -0.2 | 1.1 | 0.1 | 0.2 |
| | dATP | -0.2 | -0.2 | 0.0 | 0.2 | 0.2 | -0.1 | 0.4 | -0.2 | 0.0 |
| | dGTP | NA | NA | NA | NA | NA | NA | -0.3 | NA | 2.1 |
| | dCTP | 0.0 | -0.2 | -0.1 | 0.4 | 0.1 | 0.1 | 0.2 | 0.2 | -0.1 |
| | dUMP | -0.7 | 0.0 | 0.8 | -0.2 | -1.1 | -0.6 | -0.7 | -0.3 | -0.2 |
| | dTMP | -0.2 | 0.1 | 0.1 | 0.2 | -0.1 | 0.4 | 0.8 | -0.1 | 0.2 |
| | dTDP | 0.3 | 1.0 | 0.4 | -0.8 | -1.1 | -0.1 | 0.6 | 0.4 | 0.2 |
| | dTTP | 0.1 | 0.0 | 0.2 | 0.0 | 0.2 | -0.1 | 0.4 | 0.2 | 0.1 |
| TCA cycle and asisociated | Lac | -0.1 | NA | -0.2 | 0.2 | 0.1 | 0.4 | 0.0 | -0.1 | -0.1 |
| | Pyr | -0.3 | NA | -0.5 | 0.3 | -0.1 | 0.0 | -0.4 | -0.6 | -0.8 |
| | Cit | 0.6 | 0.0 | 0.1 | -0.2 | 0.0 | 0.0 | 0.8 | 0.2 | 0.5 |
| | ICit | 0.4 | 0.1 | -0.2 | -0.1 | 0.0 | -0.4 | 0.3 | 0.2 | 0.3 |
| | 2HG | -0.3 | -0.9 | -0.7 | 1.6 | -0.4 | -0.3 | 0.4 | -0.1 | 0.1 |
| | Suc | -1.2 | -0.8 | 0.0 | 0.5 | 0.0 | 0.5 | -0.5 | 0.3 | 0.0 |
| | Mal | -0.2 | 0.1 | -0.1 | -0.4 | -0.2 | -0.2 | 0.5 | 0.2 | 0.6 |
| Amino acids | Ala | 0.0 | NA | 0.6 | -0.2 | 0.4 | 0.2 | 0.0 | 0.5 | 0.4 |
| | Asp | 0.0 | NA | 0.1 | -0.1 | 0.0 | -0.2 | 0.0 | 0.2 | 0.0 |
| | Glu | 0.1 | NA | 0.0 | -0.1 | -0.1 | -0.2 | 0.0 | 0.0 | 0.1 |
| | Phe | 0.2 | NA | 0.5 | 0.1 | 0.5 | 0.6 | 0.0 | 0.7 | 0.7 |
| | Pro | 0.0 | NA | 0.3 | -0.1 | 0.3 | 0.1 | 0.0 | 0.2 | 0.1 |
| | Thr | 0.2 | NA | 0.5 | 0.2 | 0.6 | 0.6 | 0.1 | 0.8 | 0.7 |
| | Tyr | 0.2 | NA | 0.5 | 0.0 | 0.3 | 0.1 | 0.0 | 0.6 | 0.5 |

**Supplementary Table S1F:** Heatmapped $log_2$ fold change of all measured central carbon metabolites in Hek293 cells treated with cisplatin (1 µM), APIM-peptide (8 µM), or the respective combination for 4, 8 and 24 hours relative to untreated control cells. Blue: descreased, red: increased. Metabolite abbreviations are listed in Supplementary Table S1A.

| Class | Metabolite | 4h | | | 8h | | | 24h | | |
|---|---|---|---|---|---|---|---|---|---|---|
| | | Cisplatin | APIM-peptide | APIM-peptide + cisplatin | Cisplatin | APIM-peptide | APIM-peptide + cisplatin | Cisplatin | APIM-peptide | APIM-peptide + cisplatin |
| Glycolysis, PPP and other phosphorylated sugars | G6P | -0.2 | -0.2 | -0.7 | 0.1 | -0.1 | -0.1 | 0.0 | 0.2 | -0.1 |
| | F6P | -0.1 | -0.3 | -0.7 | 0.2 | 0.0 | 0.1 | -0.1 | 0.2 | -0.1 |
| | M6P | 0.2 | -0.3 | 0.0 | 0.0 | 0.1 | 0.0 | -0.1 | -0.4 | -0.7 |
| | G-/M1P | -0.1 | -0.1 | -0.2 | 0.1 | 0.2 | 0.1 | -0.2 | -0.2 | -0.2 |
| | F1P | -0.1 | -0.3 | -0.1 | 0.0 | 0.3 | 0.1 | -0.1 | -0.3 | -0.2 |
| | F1,6BP | 0.6 | 0.5 | 0.7 | -0.3 | -0.4 | -0.3 | 0.5 | 0.7 | 0.3 |
| | GL3P | -0.5 | -0.6 | -0.6 | -0.3 | 0.1 | 0.0 | 0.0 | 0.0 | -0.1 |
| | 2-/3PG | -0.4 | -0.3 | -0.3 | 0.5 | 0.3 | 0.3 | 0.5 | 0.4 | 0.6 |
| | PEP | -0.1 | -0.4 | -0.3 | 0.3 | 0.3 | 0.3 | 0.4 | 0.4 | 0.3 |
| | 6PG | 0.3 | 0.1 | 0.4 | -0.1 | -0.1 | -0.2 | -0.3 | 0.3 | 0.2 |
| | R5P | 0.3 | -0.1 | 0.3 | 0.5 | 0.4 | 0.1 | -0.3 | 0.1 | -0.1 |
| | PRPP | 0.2 | 0.4 | -0.4 | 1.1 | 0.7 | 0.6 | -1.2 | -1.1 | -1.9 |
| (deoxy) Nucleoside phosphates | AMP | NA | -0.4 | -1.9 | -0.7 | -0.2 | -1.2 | -1.4 | -0.7 | -0.9 |
| | ADP | -1.2 | 0.3 | -0.7 | -0.5 | -0.1 | -0.5 | -0.5 | -0.3 | -0.6 |
| | ATP | 0.0 | 0.1 | 0.1 | 0.0 | 0.0 | 0.0 | 0.0 | 0.0 | 0.0 |
| | GMP | 0.2 | 0.0 | 0.1 | 0.3 | 0.0 | 0.3 | 0.1 | -0.1 | -0.1 |
| | GDP | -0.5 | -0.2 | -0.7 | 0.2 | 0.2 | -0.4 | -0.2 | -0.5 | -0.4 |
| | GTP | 0.1 | -0.1 | -0.1 | -0.1 | -0.1 | -0.2 | 0.3 | 0.2 | 0.3 |
| | CDP | 0.0 | 0.5 | -1.8 | 0.7 | 0.7 | 0.3 | -0.6 | -0.6 | -0.9 |
| | CTP | 0.2 | -0.1 | 0.2 | 0.1 | 0.2 | 0.3 | 0.0 | 0.1 | 0.1 |
| | UMP | -0.1 | -0.2 | 0.0 | 0.1 | 0.1 | 0.1 | 0.2 | 0.2 | 0.2 |
| | UDP | -0.1 | 0.7 | -0.8 | 0.4 | 0.6 | 0.0 | -0.7 | -0.8 | -1.1 |
| | UTP | -1.2 | 0.5 | -1.1 | -0.5 | -0.1 | 1.5 | -0.1 | -0.4 | 0.1 |
| | UDP-Glc-NAc | 0.0 | -0.2 | 0.0 | 0.1 | 0.1 | 0.0 | -0.1 | -0.2 | -0.1 |
| | Citrate_GC | 0.1 | 0.0 | -0.1 | 0.0 | 0.0 | -0.1 | 0.2 | 0.2 | 0.2 |
| | D-a-OH-glutarate | 0.1 | -0.3 | 0.9 | -0.1 | 0.2 | -0.1 | -0.1 | 0.0 | 0.3 |
| | dADP | -0.3 | 0.4 | -0.8 | 0.4 | 0.4 | 0.0 | -0.5 | -1.0 | -0.7 |
| | dATP | 0.3 | 0.1 | 0.2 | 0.2 | -0.2 | 0.0 | 0.1 | -0.2 | 0.2 |
| | dGTP | NA | NA | NA | 1.1 | 1.5 | 2.1 | -2.2 | -0.4 | -0.2 |
| | dCTP | 0.1 | 0.0 | 0.1 | 0.0 | -0.2 | -0.1 | 0.1 | 0.1 | 0.2 |
| | dUMP | -0.5 | -0.4 | -0.3 | -0.5 | -0.4 | -0.3 | -0.5 | -0.1 | -0.2 |
| | dTMP | -0.1 | -0.4 | -0.2 | 0.3 | 0.0 | 0.3 | 0.0 | 0.0 | 0.4 |
| | dTDP | -0.2 | 0.6 | -0.7 | 0.8 | 0.7 | 0.2 | -0.4 | -0.6 | -0.6 |
| | dTTP | 0.0 | -0.1 | -0.1 | 0.3 | 0.0 | 0.3 | 0.2 | -0.9 | 0.1 |
| TCA cycle and asisociated | Lac | -0.1 | 0.0 | -0.1 | 0.2 | 0.0 | 0.2 | -0.1 | -0.2 | -0.2 |
| | Pyr | 0.3 | 0.1 | -0.1 | 0.0 | 0.0 | -0.1 | 0.3 | 0.2 | 0.4 |
| | Cit | -0.4 | -0.3 | -0.2 | 0.2 | 0.1 | 0.1 | 0.0 | 0.0 | 0.0 |
| | ICit | 0.1 | -0.2 | 0.1 | 0.4 | -0.1 | -0.1 | 0.0 | 0.0 | -0.1 |
| | Suc | -0.9 | -0.7 | -0.2 | 0.3 | 0.3 | 0.1 | 0.0 | 0.4 | 0.4 |
| | Fum | -0.1 | -0.1 | -0.1 | 0.0 | 0.0 | -0.1 | 0.2 | 0.2 | 0.3 |
| | Mal | -0.2 | -0.2 | 0.0 | -0.1 | 0.1 | -0.3 | -0.2 | 0.0 | 0.1 |
| Amino acids | Ala | 0.0 | 0.0 | 0.0 | 0.1 | 0.1 | 0.1 | 0.1 | 0.0 | 0.1 |
| | Asp | 0.1 | 0.0 | 0.1 | -0.3 | 0.1 | -0.3 | 0.3 | 0.3 | 0.4 |
| | Glu | 0.0 | 0.0 | 0.0 | -0.1 | 0.0 | -0.1 | 0.2 | 0.3 | 0.2 |
| | Phe | 0.0 | 0.0 | 0.1 | 0.0 | 0.1 | 0.0 | 0.1 | 0.2 | 0.2 |
| | Pro | 0.0 | -0.2 | -0.2 | 0.1 | 0.0 | 0.0 | 0.0 | 0.0 | 0.0 |
| | Thr | 0.1 | 0.1 | 0.1 | -0.1 | 0.1 | -0.1 | 0.1 | 0.2 | 0.2 |
| | Tyr | 0.0 | -0.1 | -0.1 | -0.1 | 0.0 | -0.1 | 0.1 | 0.2 | 0.2 |
| | Val | 0.0 | 0.1 | 0.1 | -0.1 | 0.1 | 0.0 | 0.1 | 0.2 | 0.2 |

**Supplementary Table S1G:** Summed fractional labeling (SFL) ± SD of selected metabolites of glycolysis, the PPP, the TCA cycle and the nucleoside phosphate pool in JJN3 cells treated with APIM-peptide (8 µM) for 4 and 24 hours relative to untreated control, n = 4. Metabolite abbreviations are listed in Supplementary Table S1A.

| Metabolite class | Metabolite | Time | Average SFL ± SD | | | | | |
|---|---|---|---|---|---|---|---|---|
| | | | 1-13$C_1$-glucose | | 1,2-13$C_2$-glucose | | U-13$C_6$-glucose | |
| | | | Ctr | APIM-peptide | Ctr | APIM-peptide | Ctr | APIM-peptide |
| Glycolysis | G6P | 4 | 17 ± 0.1 | 17 ± 0.1 | 34 ± 0.1 | 34 ± 0.4 | 93 ± 0.2 | 90 ± 1.5 |
| | | 24 | 18 ± 0.1 | 17 ± 0.4 | 35 ± 0.1 | 34 ± 0.2 | 93 ± 0.3 | 92 ± 1.6 |
| | F6P | 4 | 17 ± 0.3 | 17 ± 0.6 | 34 ± 0.2 | 33 ± 0.9 | 93 ± 0.3 | 91 ± 2.5 |
| | | 24 | 17 ± 0.3 | 17 ± 0.4 | 35 ± 0.2 | 34 ± 0.8 | 93 ± 0.7 | 91 ± 2.2 |
| | 2-/3PG | 4 | 13 ± 1.6 | 13 ± 1.1 | 31 ± 0.7 | 30 ± 1.0 | 89 ± 0.6 | 82 ± 4.5 |
| | | 24 | 13 ± 0.4 | 12 ± 0.6 | 28 ± 1.2 | 28 ± 0.9 | 88 ± 2.0 | 83 ± 7.3 |
| PPP | R5P | 4 | 12 ± 0.5 | 12 ± 0.6 | 37 ± 0.6 | 37 ± 0.6 | 93 ± 0.9 | 91 ± 1.6 |
| | | 24 | 11 ± 0.4 | 11 ± 0.4 | 36 ± 0.6 | 36 ± 0.2 | 94 ± 0.6 | 92 ± 1.6 |
| | S7P | 4 | 15 ± 0.2 | 15 ± 0.5 | 39 ± 0.1 | 38 ± 0.9 | 92 ± 0.4 | 84 ± 3.4 |
| | | 24 | 16 ± 0.1 | 15 ± 0.3 | 39 ± 0.3 | 38 ± 0.3 | 92 ± 0.6 | 87 ± 4.0 |
| TCA cycle | Cit | 4 | 8 ± 0.6 | 7 ± 3.3 | 12 ± 0.8 | 10 ± 1.3 | 22 ± 0.3 | 17 ± 1.7 |
| | | 24 | 10 ± 0.4 | 9 ± 0.6 | 17 ± 0.4 | 14 ± 0.8 | 33 ± 0.7 | 27 ± 1.4 |
| Nucleoside phosphates | ATP | 4 | 4 ± 0.0 | 4 ± 0.0 | 12 ± 0.1 | 12 ± 0.0 | 39 ± 0.0 | 38 ± 0.1 |
| | | 24 | 4 ± 0.0 | 4 ± 0.0 | 12 ± 0.1 | 12 ± 0.0 | 40 ± 0.0 | 40 ± 0.1 |
| | GTP | 4 | 7 ± 0.1 | 7 ± 0.0 | 18 ± 0.0 | 18 ± 0.1 | 44 ± 0.1 | 42 ± 0.5 |
| | | 24 | 7 ± 0.0 | 8 ± 0.1 | 19 ± 0.1 | 20 ± 0.2 | 48 ± 0.1 | 47 ± 0.4 |
| | CTP | 4 | 6 ± 0.3 | 6 ± 0.4 | 16 ± 0.1 | 15 ± 0.2 | 40 ± 0.6 | 34 ± 2.1 |
| | | 24 | 9 ± 0.2 | 8 ± 0.4 | 19 ± 0.4 | 18 ± 0.5 | 53 ± 0.4 | 50 ± 1.5 |
| | UTP | 4 | 9 ± 0.1 | 9 ± 0.2 | 22 ± 0.2 | 21 ± 0.3 | 51 ± 0.2 | 49 ± 0.6 |
| | | 24 | 11 ± 0.2 | 10 ± 0.2 | 24 ± 0.2 | 24 ± 0.6 | 58 ± 0.4 | 55 ± 1.2 |

**Supplementary Table S2:** Cell densities, stressor concentrations, replicates and sampling time pointslisted for all individual assays and cell lines/types

| Assay and cell line | Cell density (cells/ml) | Stressor concentration (µM) | | | | | | | Number of replicates | | Sampling time points (h) |
|---|---|---|---|---|---|---|---|---|---|---|---|
| | | APIM-peptide | Cisplatin | APIM-peptide + Cisplatin | mutAPIM-peptide | R11-peptide | LPS | APIM-peptide + LPS | Induividual cell cultures (n) | Repeated experiments | |
| **Viability assay (MTT)** | | | | | | | | | 6-10 | 2 | 24, 48, 72, 96 |
| JJN-3 | 50 000 | 6 | 1 | 6 + 1 | | | | | | | |
| DU145 | 15 000 | 6 | 1 | 6 + 1 | | | | | | | |
| Hek293 | 15 000 | 8 | 1 | 8 + 1 | | | | | | | |
| **Quantification of extracellular metabolites** | | | | | | | | | | | 24 |
| JJN-3 | 500 000 | 8 | 1 | 8 + 1 | 8 | 8 | | | 4-6 | 1 | |
| DU145 | 70% confluent | 8 | | | | | | | 3 | 2 | |
| Hek293 | 70% confluent | 10 | | | | | | | 3 | 2 | |
| RPMI 8226 | 500 000 | 8 | | | | | | | 4-6 | 1 | |
| MC/CAR | 500 000 | 8 | | | | | | | 4 | 3 | |
| HL60 | 500 000 | 8 | | | | | | | 4-6 | 1 | |
| NB4 | 500 000 | 8 | | | | | | | 4 | 3 | |
| **Targeted mass spectrometric metabolite profiling** | | | | | | | | | | | |
| JJN-3 | 500 000 | 8 | 1 | 8 + 1 | | | | | 3-5 (4h), 1 (8,24h) | 3 (4h), 1 (8,24h) | 4, 8, 24 |
| DU145 | 50% confluent | 8 | 1 | 8 + 1 | | | | | 1 | 1 | 4, 8, 24 |
| Hek293 | 50% confluent | 8 | 1 | 8 + 1 | | | | | 1 | 1 | 4, 8, 24 |
| RPMI 8226 | 500 000 | 8 | | | | | | | 3 | 1 | 4 |
| MC/CAR | 500 000 | 8 | | | | | | | 4 | 3 | 4 |
| HL60 | 500 000 | 8 | | | | | | | 5 | 1 | 4 |
| NB4 | 500 000 | 8 | | | | | | | 4 | 3 | 4 |
| Primary monocytes | 4 000 000* | 8 | | | | | 10 ng/ml | 8 + 10 ng/ml | 3 | 3 | 4 |
| **Multiplexed inhibitor bead (MIB)-assay** | | | | | | | | | 1 | 3 | 4 |
| JJN-3 | 150 000 | 6 | 1 | 6 + 1 | | | | | | | |
| MC/CAR | 500 000 | 8 | | | | | | | | | |
| NB4 | 500 000 | 8 | | | | | | | | | |
| Primary monocytes | 4 000 000* | 8 | | | | | 10 ng/ml | 8 + 10 ng/ml | | | |
| **Non-targeted MS profiling** | | | | | | | | | | | |
| JJN-3 | 500 000 | 8 | | 8 + 1 | | | | | 4 | 1 | 4 |
| **13C-labeling experiment** | | | | | | | | | | | |
| JJN-3 | 500 000 | 8 | | 8 + 1 | | | | | 4 | 1 | 4.24 |
| **Whole genome gene expression analysis** | | | | | | | | | | | |
| JJN-3 | 150 000 | 6 | 1 | 6 + 1 | 6 | | | | 1 | 2, 3 (24h) | 4, 8, 24 |
| **HIF1A ELISA assay** | | | | | | | | | | | |
| JJN-3 | 500 000 | 8 | | | | | | | 3** | 1 | 4, 8, 12, 24 |
| **LC-MS/MS analysis of NAD pools** | | | | | | | | | 4-6 | 1 | 4 |
| JJN-3 | 500 000 | 8 | | | 8 | 8 | | | | | |
| DU145 | 70% confluent | 8 | | | | | | | | | |
| Hek293 | 70% confluent | 8 | | | | | | | | | |
| RPMI 8226 | 500 000 | 8 | | | | | | | | | |
| HL60 | 500 000 | 8 | | | | | | | | | |
| **GSH levels** | | | | | | | | | 1*** | | 4, 24 |
| JJN-3 | 150 000 | 6 | | | | | | | | 2 | |
| DU145 | 60 000 | 6 | | | | | | | | 3 | |
| Hek293 | 60 000 | 8 | | | | | | | | 2 | |
| RPMI 8226 | 150 000 | 6 | | | | | | | | 2 | |
| HL60 | 150 000 | 6 | | | | | | | | 3 | |

*Total number of mononucleated cells seeded
**Duplicate measurements
***Triplicate measurements
